# Gap junction mediated bioelectric coordination is required for slow muscle development, organization, and function

**DOI:** 10.1101/2023.12.20.572619

**Authors:** RM Lukowicz-Bedford, JS Eisen, AC Miller

## Abstract

Bioelectrical signaling, intercellular communication facilitated by membrane potential and electrochemical coupling, is emerging as a key regulator of animal development. Gap junction (GJ) channels can mediate bioelectric signaling by creating a fast, direct pathway between cells for the movement of ions and other small molecules. In vertebrates, GJ channels are formed by a highly conserved transmembrane protein family called the Connexins. The *connexin* gene family is large and complex, presenting a challenge in identifying the specific Connexins that create channels within developing and mature tissues. Using the embryonic zebrafish neuromuscular system as a model, we identify a *connexin* conserved across vertebrate lineages, *gjd4,* which encodes the Cx46.8 protein, that mediates bioelectric signaling required for appropriate slow muscle development and function. Through a combination of mutant analysis and *in vivo* imaging we show that *gjd4*/Cx46.8 creates GJ channels specifically in developing slow muscle cells. Using genetics, pharmacology, and calcium imaging we find that spinal cord generated neural activity is transmitted to developing slow muscle cells and synchronized activity spreads via *gjd4*/Cx46.8 GJ channels. Finally, we show that bioelectrical signal propagation within the developing neuromuscular system is required for appropriate myofiber organization, and that disruption leads to defects in behavior. Our work reveals the molecular basis for GJ communication among developing muscle cells and reveals how perturbations to bioelectric signaling in the neuromuscular system may contribute to developmental myopathies. Moreover, this work underscores a critical motif of signal propagation between organ systems and highlights the pivotal role played by GJ communication in coordinating bioelectric signaling during development.

## Introduction

Animal development relies on precise coordination and communication among diverse cell types to generate complex organs and create multi-organ functional systems. Developmental decisions have been primarily understood through the lens of canonical signaling pathways and biomechanical forces. However, an equally vital mode of communication exists in the form of bioelectric signaling^1–3^. Bioelectric signaling is well-appreciated in the nervous system, where the resting membrane voltage of neurons is meticulously regulated by ion channels, pumps, and exchangers, enabling rapid transmission of information between cells^4^. Neurons share bioelectric information at either chemical or electrical synapses, with the latter utilizing gap junction (GJ) channels that enable direct and bidirectional communication of ions and small molecules^5^. Yet GJ-mediated bioelectric communication is not only used by neurons, as it is a prevalent mode of signaling across a number of tissues^6–9^. For example, GJ-synchronized calcium activity mediates contractions in cardiomyocytes maintaining consistent heartbeat rhythm^10^ and GJ-mediated oscillatory calcium patterns in pancreatic Beta-cells are necessary for glucose-stimulated insulin secretion^11^. GJ-coordinated calcium signaling instructs a diversity of developmental outcomes^12^, yet identifying the molecular basis for bioelectric signaling has been a persistent challenge for the field.

A critical barrier to understanding GJ-based cellular communication lies in the molecules that make the channels, which in vertebrates are formed by a large and highly conserved family of proteins called Connexins. Vertebrate genomes contain a diversity of Connexins, with 20 distinct Connexin-encoding genes in the human genome, and various large numbers in other species, including ∼40 in zebrafish^13^. The complexity of the Connexin family impacts GJ function as combinations of multiple Connexins can contribute to hemichannels and/or intercellular channels, influencing the permeability and functional properties of signaling^14^. Most of our knowledge of GJ channel function *in vivo* has been informed by two Connexins, Cx43, encoded by the *Gap junction alpha 1* (*Gja1*) gene, and Cx36, encoded by the *Gjd2* gene. Given the challenges of nomenclature in this gene family, we label the gene name followed by encoded protein, as this allows better tracking of homologues across phylogenies^13,15^. *Gja1*/Cx43 is broadly expressed in nonneural cells where it mediates bioelectric signaling. For example, in zebrafish fin development and regeneration, *gja1b*/Cx43 mutants have a ‘short fin’ phenotype with inappropriate proportions, underscoring the necessity for Cx43-mediated communication in appendage development and morphology^16^. Disruptions in *GJA1*/Cx43 are implicated in a diverse array of human diseases including abnormalities to the bone, skin, eye, teeth, heart, and digits, suggesting bioelectric signaling in various contexts is key to the development of a variety of developmental decisions^17,18^. By contrast, *Gjd2*/Cx36 is expressed almost exclusively in neurons where it facilitates formation of neuronal gap junctions, also called electrical synapses. In the nervous system, electrical synapses often play a critical role in synchronizing activity within populations of neurons^19–23^. Disruptions in *GJD2*/Cx36 have been implicated in a variety of neurodevelopmental disorders, including myopia, epilepsy, and autism^24,25–31^. These examples underscore the specific roles of well-known Connexins in diverse cell and tissue types. Further, they define an emerging notion of so-called “Connexinopathies”^32^, in which disruption of particular Connexin-encoding genes leads to a constellation of phenotypes based on disruption of GJ signaling. Thus, revealing the role of specific Connexins in coordinating bioelectric signaling is critical to understanding normal development and organ function and how perturbations can lead to developmental defects.

Here we focus on the role of GJ-coordinated bioelectrical communication within the developing neuromuscular system. Often, adult vertebrate skeletal muscle, including zebrafish, frog, and mammal, is typified by an absence of GJ coupling^33–36^. Yet early in development there is extensive GJ communication among developing myoblasts. For instance, developing rodent muscle cells express *Gjd4*/Cx39, *Gja1*/Cx43, and *Gjc1*/Cx45 with pervasive GJ-coupling among myocytes in early development^37–41^. During maturation, increasing neuromuscular innervation leads to decreased GJ coupling between muscle cells^33–35,42,43^. Additionally, denervating diseases such as muscular dystrophy result in aberrant muscle-muscle GJ coupling^44,45^. While GJ coupling is apparent in muscle during development and disease, mutations in muscle-expressed Connexins have not revealed definitive requirements for proper development, possibly due to redundant Connexins^38,39,46^. Using genetic approaches in zebrafish, we find that *gjd4*/Cx46.8 is specifically expressed in developing slow muscle cells where it is required to create functional GJ channels. Using calcium imaging we find that *gjd4*/Cx46.8 mediates synchronized calcium activity that is imparted to slow muscle cells via neural activity. Finally, we find that disrupting GJ communication or bioelectric signaling within the developing neuromuscular system results in defects of myosin fiber organization and behavior. These findings offer unique insights into the roles of GJ channels in coordinating bioelectric information between the developing nervous system and skeletal muscle, reveal novel insight into mechanisms underlying muscle development, and establish potential connections to myopathies.

## Results

### *gjd4* is expressed in developing slow muscle cells

During zebrafish development, GJ coupling occurs among both the slow and fast muscle fibers, but not between slow and fast fibers^43^, so we sought to comprehensively assess *connexin* expression within these cell populations. Zebrafish muscle is fast developing and accessible and by 24 hours post fertilization (hpf) functional slow and fast skeletal muscle types are readily distinguished morphologically based on position and orientation (Fig. 1A). The first skeletal muscle to develop in zebrafish is slow muscle, comprising two distinct cellular subtypes: muscle pioneers (MPs) and superficial slow fibers (SSFs)^47^. While both MPs and SSFs are born adjacent to the notochord, the numerous SSFs migrate to the surface where they are distinguished by their horizontal and parallel orientation within the somite. By contrast, the 2-6 MPs of each somite remain adjacent to the notochord^47,48^. Later-born fast muscle develops from a cell population in the posterior compartment of the somite separate from the slow muscle lineage, and also adopts two subtypes, lateral fast and medial fast fibers, with fibers oriented slightly obliquely within the somite^49^.

**Figure 1:**
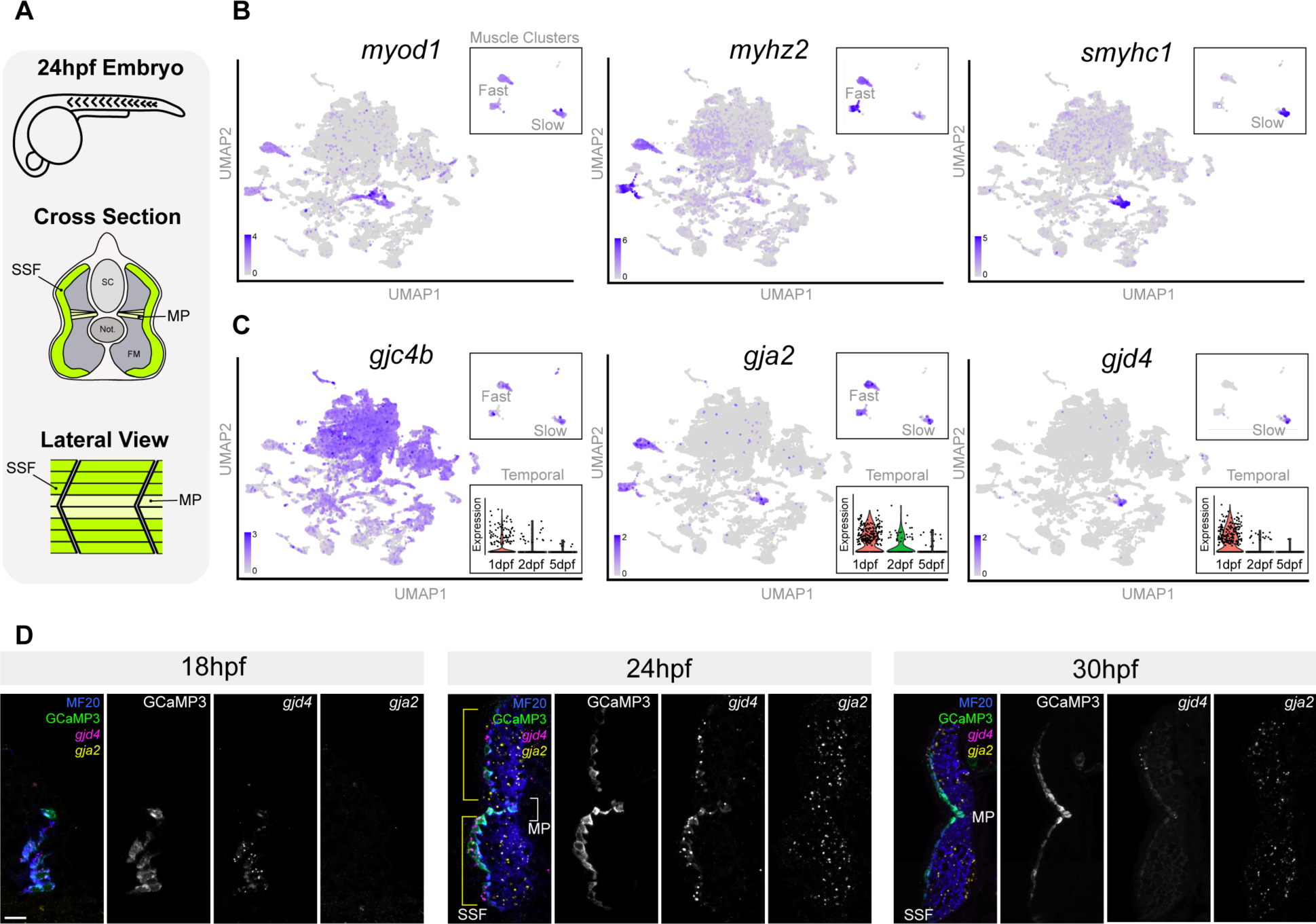
*gjd4* is expressed in developing slow muscle. **A)** Embryonic zebrafish skeletal muscle (green) is composed of superficial slow fibers (SSF) and muscle pioneers (MP). These populations are anatomically distinct: in cross-section, the SSFs reside on the most lateral surface while the MPs remain adjacent to the notochord; from a lateral view, the SSF cells lie superficially and form a chevron pattern defined by the somite boundaries, whereas the MPs reside deeper in the tissue at the dorsal/ventral midpoint of the somite. SC, spinal cord. Not., notochord. FM, fast muscle. **B)** Expression of skeletal muscle markers *myod1* (all muscle), *myzh2* (fast subtypes), and *smyhc1* (slow subtypes), are shown for all cells of the scRNAseq atlas dataset. Expression within only the putative skeletal muscle clusters is shown as subsets. **C)** Expression of three *connexin* genes, *gjc4b*, *gja2*, *gjd4*, are shown for the scRNAseq atlas and within the putative skeletal muscle clusters (upper subset). Violin plots shows expression of the specified *connexins* within the putative skeletal muscle clusters separated by the age of the cells (temporal, lower subset). **D)** Cross sections through the mid-trunk at specified times hours post fertilization (hpf), with *gjd4* (magenta) and *gja2* (yellow) labeled by fluorescent RNA in-situ hybridization, slow muscle cells visualized by *smyhc1:GCaMP3^b1464^* (green), and all muscle cells visualized by the MF20 antibody (blue). Neighboring panels show labeled individual channels in grayscale. Yellow brackets denote extent of SSF cells, white bracket denotes location of MP cells. Scale bar = 10μm.

To identify all *connexins* expressed in zebrafish muscle during development we used our single cell RNA sequencing (scRNA-seq) atlas^15,50^. In this atlas, cells were dissociated from whole embryos at 24, 48, and 120 hpf, and their individual expression profiles captured using the 10X platform. We used muscle-specific markers (*myod1, chrna1*) to identify putative skeletal muscle clusters, and type-specific markers to identify clusters related to fast (*myhz2, mylpfa*) and slow (*smyhc1, tnnc1b*) muscle subtypes (Fig. 1B, Fig. S1A)^51–53^. Within these putative slow and fast muscle clusters, we found three main *connexins* expressed at high levels in most cells of the clusters: *gjc4b, gja2*, and *gjd4* (Fig. 1C, Fig. S1B). The *gjc4b* gene encodes Cx43.4 (*gjc4b*/Cx43.4), belongs to a teleost-specific clade of *connexins*^13^, and while expressed in both developing slow and fast muscle cells, this *connexin* is notable for its nearly ubiquitous expression across cell types captured in the atlas (Fig. 1C). Previous knockdown of *gjc4b* during zebrafish development did not cause defects in gross morphological organization and function, including muscle^54^. By contrast, *gja2*/Cx39.9, another teleost-specific *connexin*^13^, is expressed specifically in developing muscle of both slow and fast types throughout the developmental timepoints captured in the atlas (Fig. 1C). Previous work found that *gja2* is expressed within slow and fast muscle, that *gja2* mutants have disrupted GJ coupling among muscle cells at 30 hpf, but that mutations in the gene do not affect the development of the tissue^46^. Finally, *gjd4*/Cx46.8 displays a restricted spatio-temporal pattern in the scRNAseq data, with exclusive expression only in developing slow muscle, and a restricted temporal window of expression at 24 hpf (Fig. 1C). The *gjd4* gene family is conserved in vertebrate lineages with zebrafish *gjd4*/Cx46.8 being the homolog of mouse *Gjd4*/Cx39 and human *GJD4*/Cx40.1^13^. In mouse, *Gjd4* is expressed during skeletal muscle embryogenesis, yet *Gjd4* mutants display no gross defects in muscle organization^38,39^. In *Gjd4* mutant mice, *Gja1*/Cx43 is upregulated, which was hypothesized to compensate for the loss of *Gjd4*/Cx39^38,39^. Taken together, the zebrafish scRNAseq data suggest there are two main *connexins*, *gjd4*/Cx46.8 and *gja2*/Cx39.9, that are specifically expressed at high levels within developing muscle.

Although the single-cell expression analysis provided insights into *connexin* expression in putative slow muscle cells, the resolution of the dataset does not allow differentiation among the distinct muscle subtypes, nor does it provide a clear temporal understanding of expression. To overcome these limitations, and to validate the scRNAseq data, we examined expression of *gjd4* and *gja2* transcripts *in vivo*. To visualize slow muscle cells, we generated a *smyhc1:GCaMP* line of transgenic fish (*Tg(smyhc1:GCaMP3^b1464^*), Fig. S2A) and detected RNA localization of *connexins* using fluorescent RNA in situ hybridization. We find that *gjd4* is expressed exclusively within developing slow muscle cells as early as 18 hpf, in both MPs and SSFs, with expression levels peaking at 24 hpf (Fig. 1D; Fig. S2B,C). We also examined *gja2* expression in the same animals and find that it is nearly undetectable at 18 hpf, is expressed in developing muscle by 24hpf, with expression persisting though diminishing as development proceeds (Fig. 1D), consistent with previous findings^46^. By analyzing *gjd4* and *gja2* expression within the same tissue, we found that *gjd4* expression precedes *gja2*, with *gjd4* being restricted to slow muscle cells, while *gja2* was predominantly expressed in fast muscle (Fig. 1D). *gja2* can be identified in slow muscle, yet it was inconsistently detected in slow muscle cells compared with fast muscle cells. Given the spatio-temporal specificity of *gjd4* expression within slow muscle cells, its conservation throughout vertebrate lineages, and its unknown function in myocytes, we focused on understanding its role in development.

### *gjd4*/Cx46.8 localizes to slow muscle cell membranes and is required for GJ communication

To investigate the role of *gjd4*/Cx46.8 in slow muscle development, we first examined localization of the *gjd4*-encoded Cx46.8 protein. We could not identify a useful antibody for this protein, thus we sought to epitope tag the Connexin in a location that would not disrupt its structure and localization. Tagging the N- or C-termini of Connexin proteins, or their transmembrane domains, disrupts processing within the ER/Golgi and/or membrane localization^55,56^, so we examined the Cx46.8 protein sequence to identify locations that might be amenable to tagging. There is a high degree of amino acid conservation among the zebrafish, mouse, and human *gjd4*-encoded Connexins within the N terminus, the four transmembrane domains, the two extracellular domains, and the intracellular loop (Fig. 2A), all regions critical to GJ structure and function^57,58^. By contrast, the intracellular C-terminal tail showed a large degree of amino acid variance among homologues (Fig. 2A), thus we targeted a non-conserved region to engineer a tag into the protein (Fig. 2B). This approach has successfully tagged a variety of proteins, including Connexins^59,60^. To visualize Cx46.8, we used CRISPR to introduce a small V5 epitope tag into the C-terminal tail of the *gjd4* locus and created a precisely edited, germline-transmitted transgenic line (Fig. 2C,D, *Pt(gjd4/Cx46.8-V5^b1459^*)). Crossing this line to the *smyhc1:GCaMP* line, we find that Cx46.8-V5 is expressed in both the MP and SSF slow muscle cell subtypes as early as 18 hpf, and protein is present until 48 hpf (Fig. 2C). Within slow muscle cells, Cx46.8-V5 expression is punctate and localizes at sites of intercellular contact, with pronounced localization at the myotendinous junctions between somites, and more dispersed localization along the dorsal and ventral surfaces of muscle cells (Fig. 2D). To visualize protein localization in individual slow muscle cells, we generated animals with mosaic Cx46.8-V5 expression, which revealed clear punctate expression on MP and SSFs, with prominent localization to the anterior and posterior membranes, and more sparse decoration of the dorsal and ventral membranes (Fig. 2E). We conclude that *gjd4*/Cx46.8 is expressed exclusively within developing MPs and SSFs, with localization patterns suggesting the protein may mediate GJ communication among cells of the developing slow muscle system.

**Figure 2:**
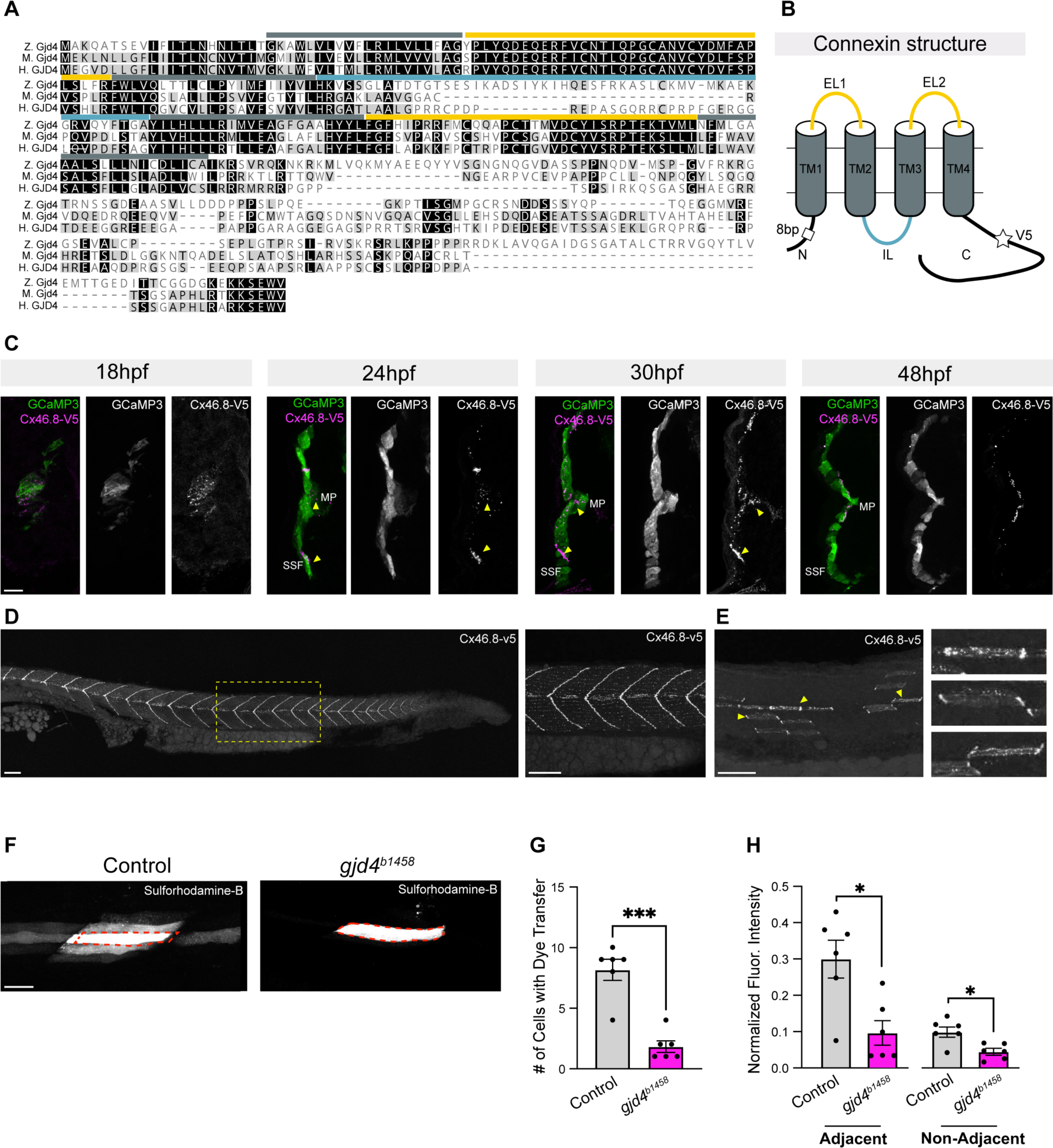
*gjd4*/Cx46.8 protein is required for GJ communication in developing slow muscle. **A)** Protein alignment of zebrafish *gjd4*/Cx46.8 with mouse Gjd4/Cx39 and human GJD4/CX40.1 orthologs. Identical amino acids are denoted with a black background and similar amino acids are denoted with a grey background. Predicted Connexin protein domains are highlighted, including transmembrane domains (TM, dark grey), extracellular loops (EL, yellow), and intracellular loop (IL, blue). **B)** Schematic of the predicted *gjd4*/Cx46.8 protein. The location of the 8bp deletion in line *gjd4^b1458^* and the V5 tag in line *Pt(gjd4/Cx46.8-V5^b1459^*) are denoted by the white box and white star, respectively. **C)** Localization of the Cx46.8-V5 protein at specified times hours post fertilization (hpf). Cross sections through the mid-trunk with Cx46.8-V5 protein visualized by the V5 antibody (magenta), and slow muscle cells visualized by *smyhc1:GCaMP3^b1464^* (green). Arrowheads note Cx46.8-V5 expression in both SSF and MP cells. Scale bar = 10μm. **D)** Lateral view of the trunk at 24 hpf showing localization of the Cx46.8-V5 protein (white). Box denotes region of zoom shown in neighboring panel to the right. Scale bar = 50μm. **E)** Mosaic, single-cell labeling of Cx46.8-V5 protein (white) at 24 hpf. Yellow arrowheads denote regions of zooms shown in neighboring panels to the right. Scale bar = 10μm. **F)** Representative images of sulforhodamine-B filled SSF cells in control and *gjd4^b1458^* mutants at 26 hpf. Scale bar = 25μm. **G)** Quantification of the number of cells with dye transfer in control and *gjd4^b1458^* mutants. Cells normalized to the filled cell, with fluorescent values of 5% or greater were included. N = 6 control and *gjd4^b1458^* mutants, unpaired t-test with Welch’s correction, p-value = 0.0002. **H)** Quantification of fluorescent intensity in cells that share a direct contact with the loaded cells (adjacent), or in cells that don’t share a direct contact with the loaded cell (non-adjacent), normalized to the loaded cell. N = 6 control and *gjd4^b1458^* mutants, unpaired t-test with Welch’s correction, adjacent cells p-value = 0.0102, non-adjacent cells p-value = 0.0110.

To assess if *gjd4*/Cx46.8 is required for slow muscle GJ coupling, we generated a CRISPR-induced 8 base pair deletion in the *gjd4* locus resulting in an early stop codon (*gjd4^b1458^*, Fig. 2B). To examine GJ coupling within developing slow muscle *in vivo* we evaluated dye transfer between cells by loading single SSFs with the GJ-permeable dye sulforhodamine-B^46^ and quantified dye fluorescence in neighboring cells relative to the dye-loaded cell. In control animals at 26 hpf, we observe robust dye transfer to adjacent SSFs, as well as to non-adjacent SSFs (Fig. 2F). In *gjd4^b1458^* mutants the number of adjacent SSFs to which dye transfers is significantly reduced, as is the extent of transfer when it occurred (Fig. 2F-H). These results are comparable to previous analyses that used pharmacological approaches to block GJ channels, resulting in loss of ionic communication and dye coupling among slow muscle cells^43,46^. Taken together, we conclude that *gjd4* encodes the Cx46.8 protein which is required to create functional GJ channels among developing slow muscle cells *in vivo*.

### *gjd4*/Cx46.8, GJ communication, and cholinergic transmission are required for coiling behavior

We examined the role of *gjd4*/Cx46.8 in slow muscle function by investigating spontaneous coiling behavior. As zebrafish develop, a series of stereotypical behaviors emerge in sequence as the neuromuscular system matures. The first behavior, termed spontaneous coiling, starts at ∼17 hpf^61–64^. To examine spontaneous coiling, we monitored spontaneous movements of control wildtype and *gjd4^b1458^* mutants at 22-23 hpf while embryos were in their chorions. Controls initiated coiling at a rate of ∼0.25 Hz, whereas *gjd4^b1458^* mutants initiated movement significantly less frequently (∼0.05 Hz, Fig. 3A). The movements of mutants in their chorions also appeared abnormal. To analyze movement kinematics, we dechorionated embryos and partially embedded them in agarose, leaving the tail free to move (Fig. 3B). Controls display typical coiling behavior with a single, smooth contraction leading to a large angle body flexure. By contrast, *gjd4^b1458^* mutants initiated movement at a lower frequency and never achieved typical coiling behavior, instead showing reduced angular distance (Fig. 3B,C). Given the specific expression of *gjd4*/Cx46.8, these findings suggest that mutants likely have disrupted slow muscle function during spontaneous coiling. Yet, we noted that animals still attempted coiling, albeit with disrupted performance.

**Figure 3:**
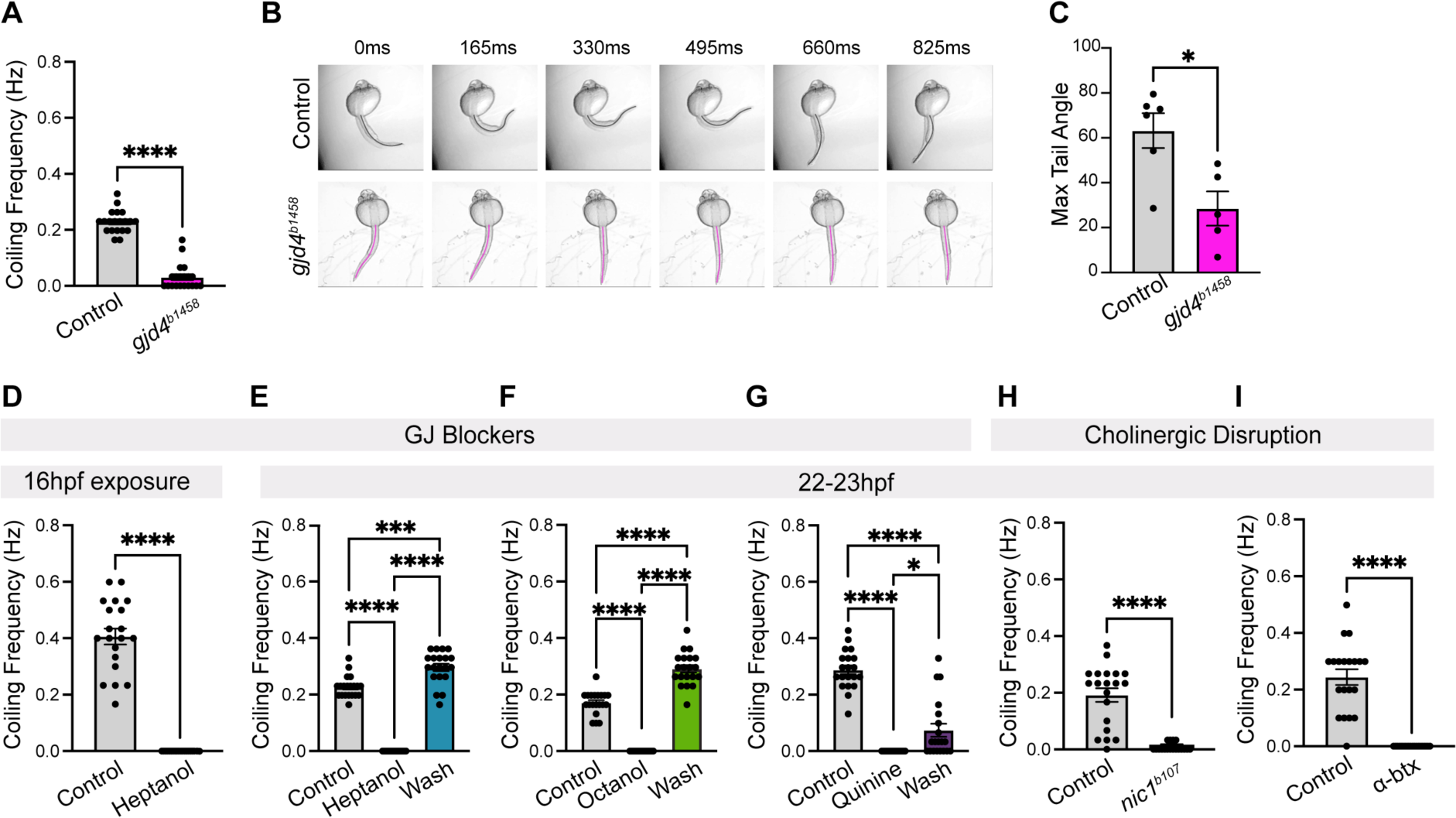
*gjd4*/Cx46.8, GJ, and cholinergic communication are required for coiling behavior. **A)** Frequency of spontaneous movement at 22-23 hpf of controls and *gjd4^b1458^* mutants. N = 20 fish per genotype, unpaired t-test with Welch’s correction, p-value = <0.0001. **B)** Representative coiling movements of control and *gjd4^b1458^* mutants. Tails are highlighted with a line for clarity in control (grey) and *gjd4^b1458^* mutants (magenta). **C)** The maximum angle achieved during coiling, measured from the tip of the tail to the yolk, in control and *gjd4^b1458^* mutants. N = 6 controls and 5 *gjd4^b1458^* mutants, unpaired t-test with Welch’s correction, p-value = 0.0293. **D-I)** Frequency of spontaneous movement at 22-23hpf. **D)** Control wildtypes and animals raised in heptanol from 16 hpf. N = 20 animals per condition, unpaired t-test with Welch’s correction, p-value = <0.0001. **E)** Animals before heptanol exposure (Control), after the animals have soaked in heptanol for 10 min (Heptanol), and after the heptanol was washed off for 10 min (Wash). N = 20 fish per condition, Brown-Forsythe ANOVA test with multiple comparisons, pre-heptanol vs. heptanol p-value = <0.0001, pre-heptanol vs. wash p-value = 0.0003 and heptanol vs. wash p-value = <0.0001. **F)** Animals before octanol exposure (Pre), after the animals have soaked in octanol for 10 min (Octanol), and after the octanol was washed off for 10 min (Wash). N = 20 animals per condition, Brown-Forsythe ANOVA test with multiple comparisons was run, pre-octanol vs. octanol p-value = <0.0001, pre-octanol vs. wash p-value = <0.0001, and octanol vs. wash p-value = <0.0001. **G)** Animals before quinine exposure (Control), after the animals have soaked in quinine for 10 min (Quinine), and after the quinine was washed off for 10 min (Wash). N = 20 animals per condition, Brown-Forsythe ANOVA test with multiple comparisons, pre-quinine vs. quinine p-value = <0.0001, pre-quinine vs. wash p-value = <0.0001, and quinine vs. wash p-value = 0.0133. **H)** Control wildtypes and *nic1^b107^* mutants. N = 20 animals per condition, unpaired t-test with Welch’s correction, p-value = <0.0001. **I)** Control, water sham-injected, animals and alpha-bungarotoxin (α-btx) injected animals. Injections occurred at 16 hpf. N = 20 fish per injection, unpaired t-test with Welch’s correction, p-value = <0.0001.

We hypothesized *gjd4*/Cx46.8-independent GJ coupling might explain the residual behavior in *gjd4^b1458^* mutants. Coiling behavior relies on two separate GJ-coupled systems, neuronal and muscular, connected to one another via cholinergic chemical synapses at the neuromuscular junction (NMJ). The spontaneous neural network activity driving coiling originates from pacemaker cells that initiate periodic depolarizations among GJ-coupled spinal cord circuitry^23,61,65–68^. Primary motoneurons (PMNs) relay neural circuit activity to developing slow muscle cells, with coiling beginning shortly after the first NMJ synapses form^48,62,69,70^. Cholinergic input from PMNs creates postsynaptic currents that spread among GJ-coupled slow muscle cells to drive coordinated coiling behavior^43^. To test the hypothesis that *gjd4*/Cx46.8 contributes to only a subset of GJ function within the developing neuromuscular system, we turned to pharmacological and genetic approaches to broadly inhibit GJ channels and cholinergic communication. We first used heptanol, a compound that inhibits GJ communication in both neurons and muscle cells at these early timepoints in development^23,46,71^. Adding heptanol to embryos before the onset of coiling at ∼16 hpf results in a complete absence of coiling at 22-23 hpf (Fig. 3D). We further find that acute application of heptanol at 22-23 hpf resulted in a complete cessation of coiling within 10 minutes, an effect that was reversed by washing heptanol out of the medium (Fig. 3E). Given that heptanol also has non-GJ effects on cellular physiology^72^, we used the alternative GJ blockers octanol and quinine, as each has distinct off-target effects^73^. In all cases, acute application of GJ blockers results in a complete loss of coiling behavior that was reversible upon agent washout (Fig. 3E-G). These results support a model in which slow muscle GJ coupling mediated by *gjd4*/Cx46.8 is part of a larger GJ-coupled network necessary for appropriate coiling behavior.

To isolate the muscular system from the neuronal, we tested whether cholinergic communication from motoneurons was required for coiling by disrupting synaptic communication to developing muscle. We first used *nic1^b107^* mutants in which the alpha subunit of nicotinic muscle acetyl choline receptors (AChRs) is disrupted, preventing their functional assembly^74^. In *nic1^b107^* mutants, coiling movements are not observed at 17 hpf (Fig. 3H), and animals are paralyzed through 25 hpf, consistent with previous results^75^. Additionally, the application of alpha-bungarotoxin (α-btx), which blocks neuromuscular junction cholinergic receptors^76^, results in a lack of coiling behavior (Fig. 3I). Taken together, these findings indicate that *gjd4*/Cx46.8 is essential for the appropriate performance of spontaneous coiling, yet suggests it plays a specific role in coordinating bioelectric signaling within the developing neuromuscular system.

### *gjd4*/Cx46.8 is required for synchronized calcium dynamics in SSFs

Given that *gjd4*/Cx46.8 is expressed in slow muscle, we hypothesized that the behavioral deficits in mutants result from defects in the ability to coordinate muscle contractions. Cholinergic activation of slow muscle elicits ryanodine-receptor-mediated intracellular calcium release to activate muscle contraction^43,75,77,78^, so we explored if *gjd4*/Cx46.8 affects calcium activity within developing slow muscle cells. We used the *smyhc1:GCaMP^b1464^* line and imaged 3 somites at the posterior end of the yolk extension (somites 15-18) for 330 seconds at 23-24 hpf. As coiling relies on slow muscle calcium activity, we did not use pharmacological manipulations to immobilize animal, instead we achieved stable imaging by embedding them in agar which enabled us to track calcium dynamics in individual cells. Examination of the superficially located SSF slow muscle cells in control animals reveals rhythmic, coordinated calcium bursting, with tight correlation among individual cells across the field of view (Fig. 4A). Cell-to-cell correlations are tightly coupled regardless of where the cells are located relative to one another (Fig. S3A). By contrast, in *gjd4^b1458^* mutants, SSF cells show almost no dynamic calcium activity, and the remaining fluctuations lack apparent correlation between cells (Fig. 4B). While most *gjd4^b1458^* mutants exhibit a near complete loss of calcium activity in SSFs, there are occasional ‘bursting’ cells with uncorrelated sporadic activity (e.g., Fish 3 in Fig. 4B). Through cross-correlation analysis, we find that *gjd4^b1458^* mutants exhibit a significant reduction of correlated activity among SSF cells (Fig. 4C,C’). Furthermore, the event intensity and frequency of calcium activity is significantly altered in mutants (Fig. 4D-E). We conclude that *gjd4*/Cx46.8 is required for coordinated calcium dynamics among developing SSF cells.

**Figure 4:**
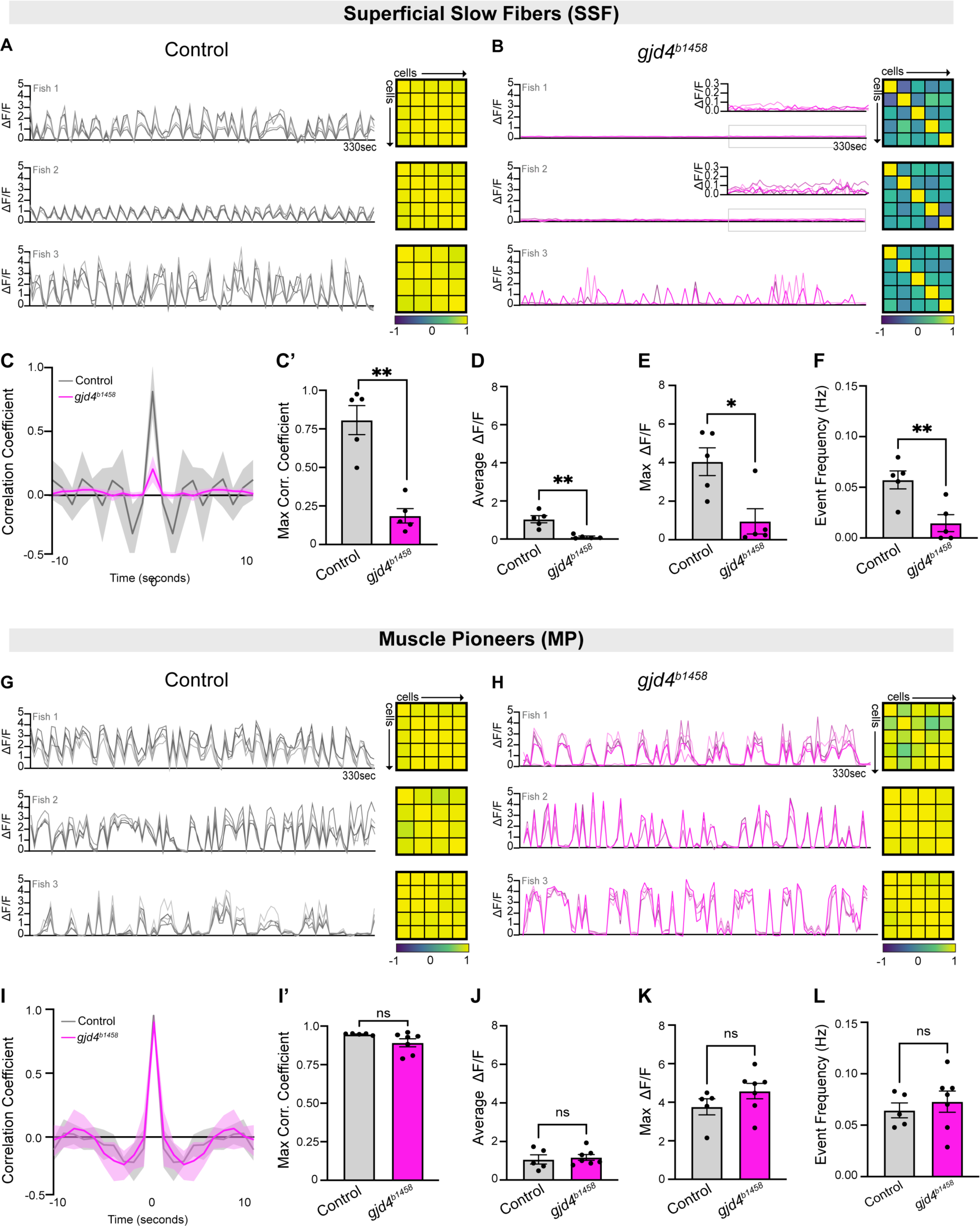
*gjd4/*Cx46.8 is required for synchronized calcium activity in SSF cells. **A-F)** Analysis of calcium dynamics in SSF cells. **A)** Control SSF GCaMP fluorescence traces of individual cells from three animals (Fish 1-3) over 330 seconds; each cell is shown as a different shade of grey. Pearson’s correlation matrix for each animal is shown to the right, yellow represents positive correlation values, and purple represents negative correlation values. **B)** *gjd4^b1458^* mutant SSF GCaMP fluorescence traces of individual cells from three different animals over 330 seconds, each cell is shown as a different shade of pink. Pearson’s correlation matrix for each animal is shown to the right. **C)** Cross-correlation analysis showing mean (dark line) and standard deviation (shading) of control (grey) and *gjd4^b1458^* mutant (magenta) SSF cells. N = 5 animals per genotype. **C’)** Max correlation coefficient for each animal. N = 5 animals per genotype, unpaired t-test with Welch’s correction, p-value = 0.0011. **D)** Average GCaMP fluorescence across the recording session for each animal. N = 5 animals per genotype, unpaired t-test with Welch’s correction, p-value = 0.0115. **E)** Max GCaMP fluorescence across the recording session for each animal. N = 5 animals per genotype, unpaired t-test with Welch’s correction, p-value =0.0132. **F)** Average peak frequency across the recording session for each animal. N = 5 animals per genotype, unpaired t-test with Welch’s correction, p-value = 0.0081. G-L) Analysis of calcium dynamics in MP cells. **G)** Control MP GCaMP fluorescence traces of individual cells from three different animals over 330 seconds, each cell is shown as a different shade of grey. Pearson’s correlation matrix for each animal is shown to the right. **H)** *gjd4^b1458^* mutant MP GCaMP fluorescence traces of individual cells from three different animals over 330 seconds, each cell is shown as a different shade of pink. Pearson’s correlation matrix for each animal is shown to the right. **I)** Cross-correlation analysis showing mean (dark line) and standard deviation (shading) of control (grey) and *gjd4^b1458^* mutant (magenta) MP cells. N = 5 controls and 7 mutants. **I’)** Max correlation coefficient for each animal. N = 5 controls and 7 mutants. Unpaired t-test with Welch’s correction, p-value = 0.0780. **J)** Average GCaMP fluorescence across the recording session for each animal. N = 5 controls and 7 mutants, unpaired t-test with Welch’s correction, p-value = 0.7191. **K)** Max GCaMP fluorescence across the recording session for each animal. N = 5 controls and 7 mutants, unpaired t-test with Welch’s correction, p-value = 0.1913. **L)** Average peak frequency across the recording session for each animal. N = 5 controls and 7 mutants, unpaired t-test with Welch’s correction, p-value = 0.5171.

After finding that *gjd4*/Cx46.8 is required for coordinated calcium activity in SSFs, we wanted to understand whether *gjd4*/Cx46.8 expression in MPs contributes to similar dynamics. Given the deeper location of the cells, we examined MP calcium activity in separate imaging experiments. In control animals, MPs have coordinated calcium dynamics with a strong correlation amongst cells within and between somites, and the dynamic characteristics are comparable in frequency and amplitude to those observed in control SSFs (Fig. 4G, Fig. S3B-C). In *gjd4^b1458^* mutants, MPs exhibit calcium dynamics that appear unaffected compared to control MP activity, displaying rhythmic and highly correlated activity among MP cells (Fig. 4H). Cross-correlation analysis and descriptive properties of dynamics do not show significant changes in *gjd4^b1458^* mutant MPs (Fig. 4I-L). Thus, despite *gjd4*/Cx46.8 being expressed in both MPs and SSFs, we conclude it is critical for coordinated calcium dynamics only among developing SSF cells, and that dynamic and correlated calcium changes in MPs can occur independent of these channels.

### GJ communication and cholinergic transmission are required for spontaneous calcium activity in developing slow muscle cells

Given that we observed a complete loss of behavior when using GJ blocking agents (Fig. 3D-G), we reasoned that we could inhibit both MP and SSF calcium dynamics by globally inhibiting GJs. To test this notion, we acutely added GJ blockers at 23 hpf and imaged SSF and MP calcium dynamics in the *smyhc1:GCaMP ^b1464^* line (Fig. 5). When treated with the GJ blocker heptanol for 10 minutes, SSFs show reduced correlated activity that is restored to control levels by washout (Fig. 5A-D). In contrast to *gjd4^b1458^* mutants, MPs in heptanol-treated animals become uncorrelated, having only rare and infrequent calcium dynamics that recover after washout (Fig. 5I-L). We find a similar loss of activity with treatment using the GJ blocker octanol, with washout restoring coordinated calcium dynamics (Fig. S4). In contrast to the specific requirement for *gjd4*/Cx46.8 in SSF cell calcium dynamics, these results support the notion that GJ channels throughout the developing neuromuscular system are required for coordinated activity in both MP and SSF subtypes.

**Figure 5:**
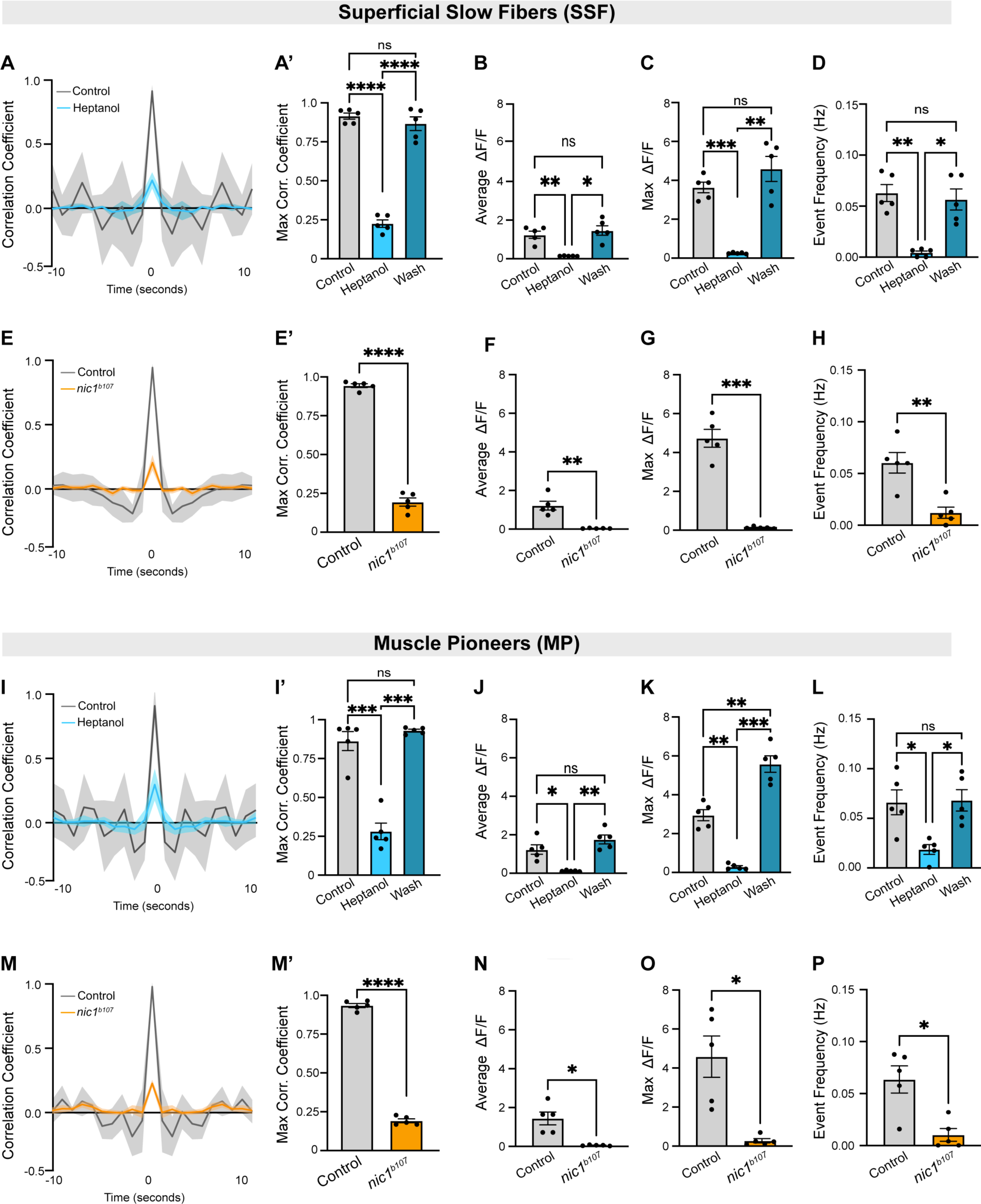
Gap junction and cholinergic communication are required for spontaneous calcium activity in slow muscle cells. **A-H)** Analysis of calcium dynamics in SSF cells. **A)** Cross-correlation analysis of SSF cells showing mean (dark line) and standard deviation (shading) of control (grey) and heptanol treated (blue) embryos. N = 5 animals per condition. Heptanol-wash cross-correlation traces were omitted for graphical clarity. A’) Max correlation coefficient for each animal, Brown-Forsythe ANOVA test with multiple comparisons, control vs. heptanol p-value = <0.0001, control vs. wash p-value = 0.6918, heptanol vs. wash p-value = <0.0001. **B)** Average GCaMP fluorescence across the recording session for each animal. N = 5 animals per condition, Brown-Forsythe ANOVA test with multiple comparisons, control vs. heptanol p-value = 0.0098, control vs. wash p-value = 0.8534, and heptanol vs. wash p-value = 0.0141. **C)** Max GCaMP fluorescence across the recording session for each animal. N = 5 animals per condition, Brown-Forsythe ANOVA test with multiple comparisons, control vs. heptanol p-value = 0.0006, control vs. wash p-value = 0.4953, and heptanol vs. wash p-value = 0.0065. **D)** Average peak frequency across the recording session for each animal. N = 5 animals per condition, Brown-Forsythe ANOVA test with multiple comparisons, control vs. heptanol p-value = 0.0059, control vs. wash p-value = 0.9478 and heptanol vs. wash p-value = 0.0190. **E)** Cross-correlation analysis of SSF cells showing mean (dark line) and standard deviation (shading) of control (grey) and *nic1^b107^* mutant (orange) animals. N = 5 animals per genotype. **E’)** Max correlation coefficient for each animal. N = 5 animals per genotype, unpaired t-test with Welch’s correction, p-value = <0.0001. **F)** Average GCaMP fluorescence across the recording session for each animal. N = 5 animals per genotype, unpaired t-test with Welch’s correction, p-value = 0.0062. **G)** Max GCaMP fluorescence across the recording session for each animal N = 5 animals per genotype, unpaired t-test with Welch’s correction, p-value = 0.0006. **H)** Average peak frequency across the recording session for each animal N = 5 animals per genotype, unpaired t-test with Welch’s correction, p-value = 0.0049. I-P) Analysis of calcium dynamics in MP cells. **I)** Cross-correlation analysis of MP cells showing mean (dark line) and standard deviation (shading) of control (grey) heptanol treated (blue) embryos. N = 5 animals per condition. Heptanol-wash cross-correlation traces were omitted for graphical clarity. **I’)** Max correlation coefficient for each animal, Brown-Forsythe ANOVA test with multiple comparisons, control vs. heptanol p-value = 0.0003, control vs. wash p=value = 0.6530, heptanol vs. wash p-value = 0.0007. **J)** Average GCaMP fluorescence across the recording session for each animal. N = 5 animals per condition, Brown-Forsythe ANOVA test with multiple comparisons, control vs. heptanol p-value = 0.0256, control vs. wash p-value = 0.3707 and heptanol vs. wash p-value = 0.0050. **K)** Max GCaMP fluorescence across the recording session for each animal. N = 5 animals per condition, Brown-Forsythe ANOVA test with multiple comparisons, control vs. heptanol p-value = 0.0021, control vs. wash p-value = 0.0038 and heptanol vs. wash p-value = 0.0006. **L)** Average peak frequency across the recording session for each animal. N = 5 animals per condition, Brown-Forsythe ANOVA test with multiple comparisons, control vs. heptanol p-value = 0.0431, control vs. wash p-value = 0.9991 and heptanol vs. wash p-value = 0.0162. **M)** Cross-correlation analysis of MP cells showing mean (dark line) and standard deviation (shading) of control (grey) and *nic1^b107^* mutant (orange) animals. N = 5 animals per genotype. **M’)** Max correlation coefficient for each animal, unpaired t-test with Welch’s correction, p-value = <0.0001. **N)** Average GCaMP fluorescence across the recording session for each animal. N = 5 animals per genotype, unpaired t-test with Welch’s correction, p-value = 0.0136. **O)** Max GCaMP fluorescence across the recording session for each animal. N = 5 animals per genotype, unpaired t-test with Welch’s correction, p-value = 0.0149. **P)** Average peak frequency across the recording session for each animal. N = 5 animals per genotype, unpaired t-test with Welch’s correction, p-value = 0.0113.

To examine whether the calcium dynamics we observed in *gjd4^b1458^* mutant MPs are driven by cholinergic transmission from motoneurons, we imaged calcium dynamics in SSFs and MPs of *nic1^b107^* mutants at 23-24hpf. In *nic1^b107^* mutants, there is a nearly complete loss of dynamic calcium activity and a significant reduction in correlation among SSFs (Fig. 5E-H) and MPs (Fig. 5M-P). We observe a similar loss of calcium dynamics in both SSFs and MPs in animals injected with alpha-bungarotoxin (α-btx) (Fig. S4). These results are in line with previous work showing that preventing cholinergic transmission in zebrafish embryos inhibits calcium dynamics in muscle^75^. Our results reveal that both MP and SSF slow muscle subtypes are affected. Taken together, results from our calcium imaging experiments support the notion that bioelectrical signaling in the embryonic neuromuscular system relies on GJ-mediated dynamic activity in neurons, communicated to developing MPs via cholinergic transmission from PMNs, and that GJs constructed from *gjd4*/Cx46.8 coordinate calcium dynamics among developing slow muscle cells.

### *gjd4*/Cx46.8 and bioelectric activity are required for slow muscle structural organization

Coordinated calcium dynamics mediated by GJ channels are critical for developmental outcomes in a variety of tissues^12,78,79^, so we next sought to investigate the role of *gjd4*/Cx46.8 in slow muscle patterning and morphology. Based on the specific expression and function *of gjd4*/Cx46.8 within slow muscle, we first focused on the structural organization of myofibers in *gjd4^b1458^* mutants. Slow muscle myosin staining reveals significant disruptions to the organization of slow muscle fibers in 24 hpf *gjd4^b1458^* mutants. *gjd4^b1458^* mutant SSFs are thinner than those of wildtypes and appear wrinkled, which we term ‘wispy and crinkly’ (Fig. 6A). Individual fibers exhibit increased tortuosity (Fig. 6B,C) and had shorter sarcomeres (Fig. 6B,D), a phenotype that persists up to 48 hpf (Fig. S6A-C). By contrast, the morphology, tortuosity, and sarcomere size of *gjd4^b1458^* mutant MPs are not significantly different from controls (Fig. 6A-D).

**Figure 6:**
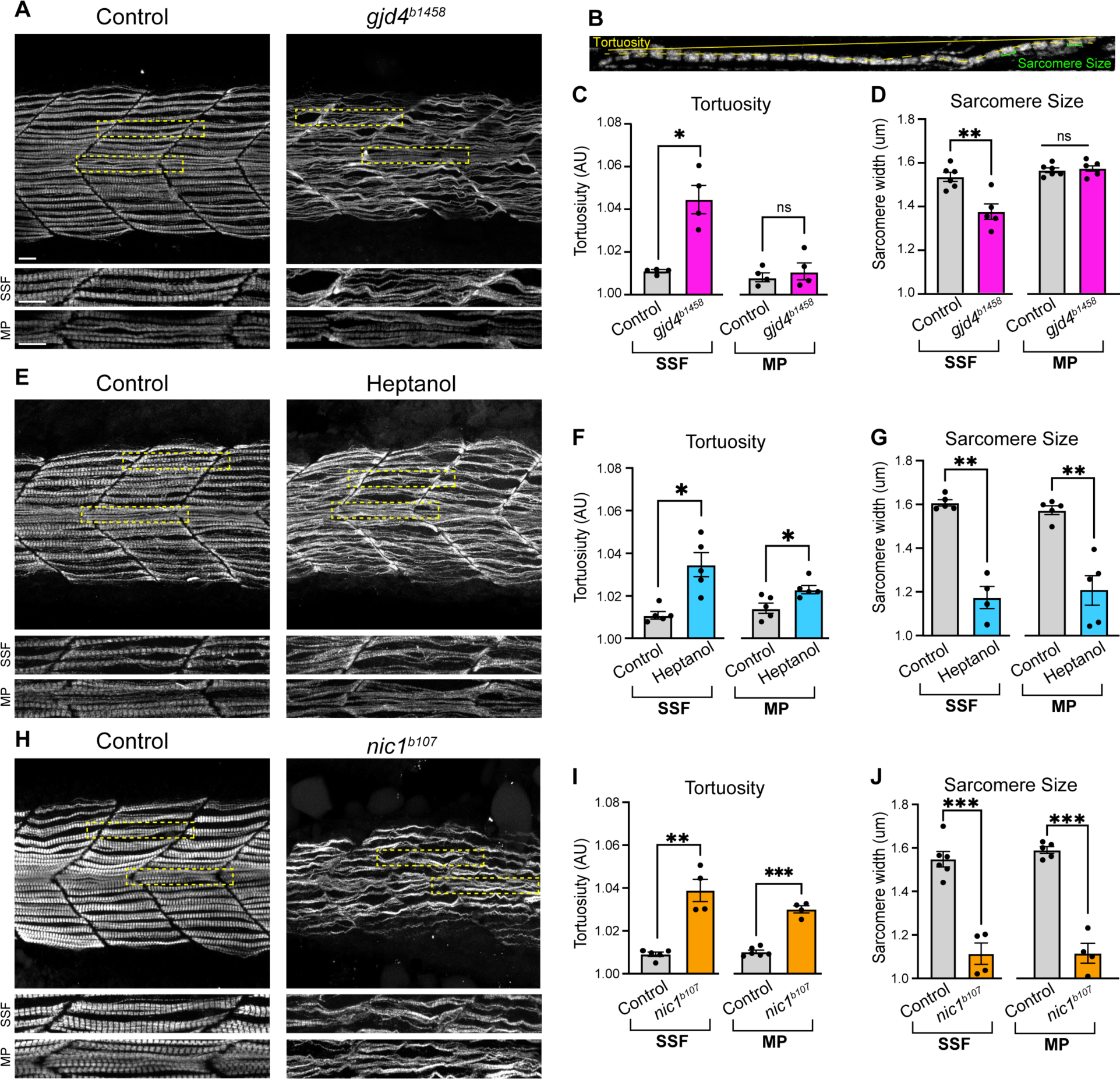
*gjd4*/Cx46.8 and bioelectric activity are required for slow muscle structural organization. **A)** Representative image of slow muscle myosin fibers at 24 hpf in control and *gjd4^b1458^* mutant animals. Yellow-dashed boxes indicate zoomed regions for the SSF or MP, shown in neighboring panels below. Scale bar = 10μm. **B)** Representation of tortuosity, measured as the difference of fiber length and somite length (yellow), and sarcomere size, measured as sarcomere width (green). **C)** Quantification of tortuosity of SSF and MP cells in controls or *gjd4^b1458^* mutants. N = 4 animals per genotype, unpaired t-test with Welch’s correction. SSF p-value = 0.0143. MP p-value = 0.5620. **D)** Quantification of sarcomere size of SSF and MP cells in control or *gjd4^b1458^* mutants. N = 6 control, 5 *gjd4^b1458^* mutants, unpaired t-test with Welch’s correction. SSF p-value = 0.0070. MP p-value = 0.8535. **E)** Representative images of slow muscle myosin fibers at 24 hpf in control or heptanol treated animals. Yellow-dashed boxes indicate zoomed regions for SSF or MP, shown in neighboring panels below. Scale bar = 10μm. **F)** Quantification of the tortuosity of SSF and MP cells in controls or heptanol treated animals. N = 5 animals per condition, unpaired t-test with Welch’s correction. SSF p-value = 0.0114. MP p-value = 0.0229. **G)** Quantification of sarcomere size of SSF and MP cells in controls or heptanol treated animals. For SSF, N = 5 controls and 4 heptanol treated animals, for MP, N = 5 animals per condition, unpaired t-test with Welch’s correction SSF p-value = 0.0021 and MP p-value = 0.0041. **H)** Representative image of slow muscle myosin fibers at 24 hpf in a control or *nic1^b107^* mutants. Yellow-dashed boxes indicate zoomed regions for SSF or MP, shown in neighboring panels below. Scale bar = 10μm. **I)** Quantification of tortuosity of SSF and MP cells in controls or *nic1^b107^* mutants. N = 5 control animals and 4 *nic1^b107^* mutants, unpaired t-test with Welch’s correction. SSF p-value = 0.0092. MP p-value = 0.0002. **J)** Quantification of sarcomere size of SSF and MP cells in controls or *nic1^b107^* mutants. For SSF, N = 6 control and 4 *nic1^b107^* mutants, for MP, N = 5 control animals and 4 *nic1^b107^* mutants, unpaired t-test with Welch’s correction for SSF p-value = 0.0004 and MP p-value = 0.0009.

We also examined whether there are additional developmental defects in *gjd4^b1458^* mutants. Previous work found that slow muscle subtype specification relies on sonic hedgehog (Shh) signaling which requires calcium release from ryanodine receptors for appropriate signaling^77^. Given the calcium defects in *gjd4^b1458^* mutants, we examined Shh signaling using *Tg(ptch2:kaede^a4596^*) that acts as a transcriptional reporter of the pathway^80,81^. In contrast to the disruption of myosin fiber organization, we find that *ptch2* expression is unaffected in *gjd4^b1458^* mutants (Fig. S5A). Additionally, using Prox1 and Engrailed transcription factor labeling^77^., we find no significant changes in the number of MP, SSF, or MFF muscle subtypes between controls and *gjd4^b1458^* mutants (Fig. S5B). Finally, we examined various features of muscle morphology, including the myoseptum (laminin), dystrophin, and fast muscle, and find no gross disruptions in mutants compared to controls (Fig. S5C). Overall, these findings suggest that the *gjd4^b1458^* mutation disrupts myosin fiber organization in SSFs but does not affect gross aspects of neuromuscular development within somites.

After finding that GJ channels, cholinergic transmission, and calcium dynamics are required in both MPs and SSFs for coiling behavior, we examined the effects of disrupting these signaling systems on the myosin organization of slow muscle fibers. We inhibited GJ channels or cholinergic transmission and observed myosin structure of slow muscle cells at 24 hpf. SSFs of animals treated with the GJ blocker heptanol display defective structural organization, increased tortuosity, and reduced sarcomere size (Fig. 6E-G). In contrast to *gjd4^b1458^*, MPs of heptanol treated animals display defects in morphology and sarcomere size (Fig. 6E-G). These phenotypes are recapitulated by other GJ blockers (Fig. S6D-F). Blocking cholinergic transmission via *nic1^b107^* mutants also affects both MPs and SSFs morphology, tortuosity, and sarcomere size (Fig. 6H-J). These phenotypes are recapitulated through inhibiting cholinergic communication via α-btx (Fig. S6G-I). These findings strongly support the notion that GJ-mediated bioelectric signaling in the developing neuromuscular system, cholinergic input from PMNs to MPs, and *gjd4*/Cx46.8 GJ channels are required for normal muscle morphology, sarcomere size, and overall fiber organization and development.

## Discussion

In this study we characterize a conserved connexin, *gjd4*/Cx46.8, that mediates bioelectric signaling during neuromuscular development in zebrafish. We reveal that *gjd4*/Cx46.8 has a specific role in developing slow muscle where it is necessary for GJ communication and coordinated calcium activity among the SSF cell subtype. Using genetic and pharmacological approaches, we find that a larger GJ coupled network is required for activity throughout the developing neuromuscular system and disruption of GJ activity or cholinergic transmission from motoneurons affects dynamic activity in both MP and SSF slow muscle subtypes. Taken together, our results support a model in which bioelectric neural activity is initiated by GJ-coupled spinal cord networks that specifically activate MPs via cholinergic input from PMNs, with coordinated activity being propagated among SSFs via *gjd4*/Cx46.8-GJ channels (Fig. 7A). Strikingly, disrupting dynamic activity in the developing system leads to defects in the structural organization of slow muscle fibers and ultimately to defects in behavior (Fig. 7B). This work delineates how bioelectric information originating in one system, here the nervous system, can influence another, skeletal muscle, and highlights how GJ channels are integral to coordination of activity dynamics and functional developmental outcomes.

**Figure 7:**
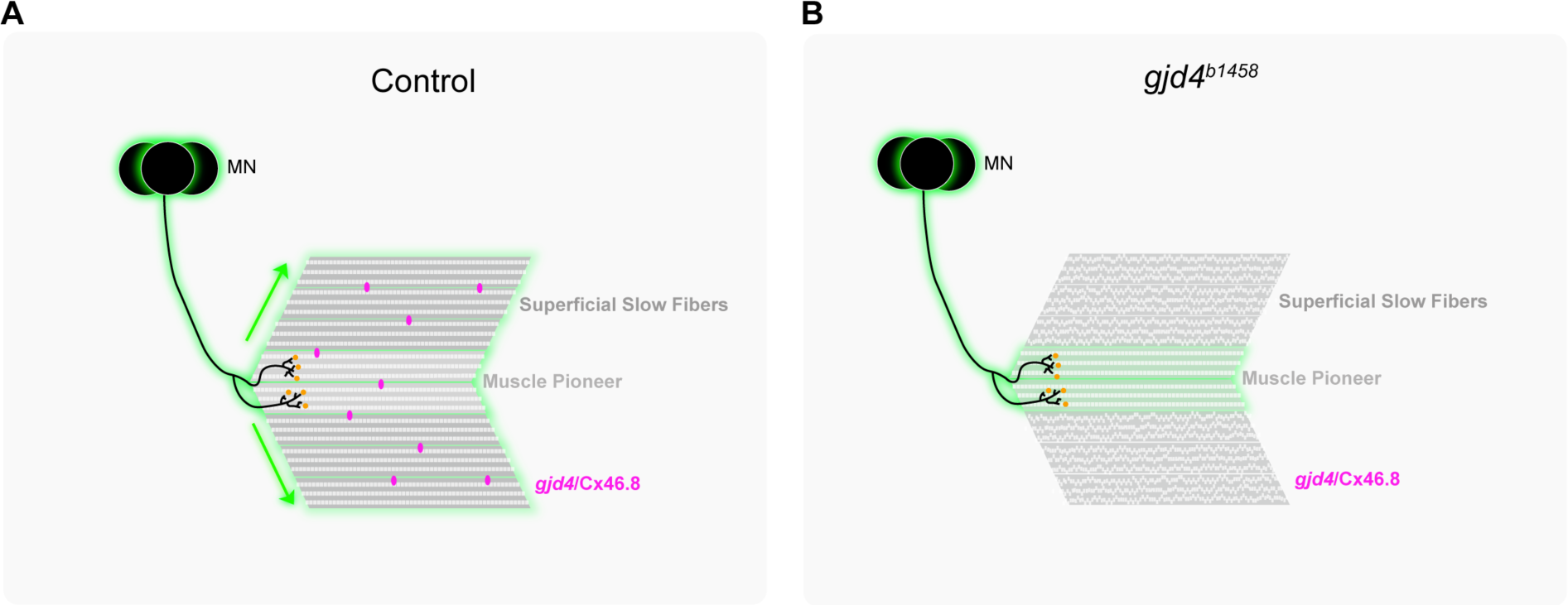
*gjd4*/Cx46.8 is required for gap junction mediated bioelectric coordination of slow muscle development, organization, and function. A) Model in which coordinated bioelectric information from the nervous system activates MPs through cholinergic synapses (orange), and the MPs propagate that synchronized information (green) to the remaining SSF via *gjd4*/Cx46.8 gap junction channels (magenta). Propagation of bioelectric information is required for proper slow myosin fiber organization and development. **B)** Without *gjd4*/Cx46.8, the nervous system still activates MPs, but the signal is unable to propagate to the remaining SSF cells, resulting in frayed myosin fibers and smaller sarcomeres. Not depicted – when GJs are disrupted globally, or when cholinergic synapses are inhibited, myosin fiber organization and development are disrupted in both MP and SSF cells.

The presence of GJ channels among myocytes in early development has long been appreciated, yet the complexity of the *connexin* gene family has created challenges in identifying specific roles for individual genes. While we focused on *gjd4*/Cx46.8 here, we found that other Connexins are also present, including *gja2*/Cx39.9, which is expressed later in development^46^. Mutations in *gja2*/Cx39.9 have defects in GJ coupling among muscle cells after the time points we investigated, resulting in behavioral defects in touch-evoked coiling which emerges after spontaneous coiling. However, *gja2*/Cx39.9 mutants do not display defects in muscle development^46^, which contrasts with our findings for *gjd4*/Cx46.8 mutants. How do the complexities of Connexin usage play out in muscle cells? The expression and mutational analyses of these two *connexins* suggest that *gjd4*/Cx46.8 may play a pioneering role during initial muscle development, while *gja2*/Cx39.9 becomes important as the system matures. Additionally, it is possible that *gjd4*/Cx46.8 and *gja2*/Cx39.9 function in concert to mediate muscle development. In other systems, Connexins can contribute to the same GJ structure by being found adjacent to one another at the same sites of cellular contact or by creating complex hemichannels and/or channels composed of multiple distinct Connexins^14,82–84^. These complexities of Connexin usage may explain why *Gjd4-/-* mice do not display overt defects in muscle development – indeed, *Gja1*/Cx43 is upregulated in these mouse mutants and may rescue normal function^38,39^. It is tempting to speculate that tackling the complexity of Connexin expression in embryonic mammalian muscle would reveal requirements similar to those we discovered in zebrafish for GJ coupling in neuromuscular development.

Our study also highlights the role of GJ channel communication in orchestrating the transmission of bioelectric information among muscle cells, concurrent with the ongoing innervation of muscle by motoneurons. The relationship between motoneuron innervation and the presence of GJ channels in skeletal muscles is critical. First, as the neuromuscular system develops and myofibers become innervated by motoneurons, there is concurrent loss of GJ coupling among muscle cells^33–35,42,43^. Consistent with this, prolonged GJ coupling is found in frog skeletal muscle upon blocking cholinergic transmission^33^ and abnormal expression of Connexins is observed in muscular dystrophy patients with denervation^44,45^. It is also noteworthy that skeletal muscles of many vertebrates are one of the few cell types that do not typically display GJ-mediated communication in their adult or mature stages^35^. Thus, we hypothesize that embryonic skeletal muscles rely on GJ communication to propagate bioelectric signals from the nervous system, but as motoneurons mature and establish innervation, the necessity for GJ-mediated communication diminishes. While not directly tested in our experiments, the disruption to sarcomere size and myosin fibers in our drug exposed and mutant fish suggests these processes are activity dependent. Our results establish the critical role of early GJ coupling as required for appropriate muscle development, and this suggests that perturbations to bioelectric signaling in early neuromuscular development could have important implications for developmental myopathies.

Our research has centered on investigating the involvement of GJ channels in transmission of bioelectric information during neuromuscular development. However, we propose that GJ-mediated bioelectric coordination between systems is likely to be a broad functional motif across various cell types in multicellular development. An intriguing example that parallels our findings is observed in the pancreas. Pancreatic cells, including beta cells, are innervated and activated by the vagus nerve^85^ and exhibit GJ-channel mediated oscillatory and rhythmic calcium activity that is necessary for glucose-stimulated insulin secretion^11^. These and other examples^1–3^ make it enticing to speculate that GJ-coordinated bioelectric signaling between organ systems will determine a wide array of development outcomes. Yet, given the large family of Connexin-encoding genes, the specific *connexins* that mediate bioelectric dynamics across developing multicellular systems remain largely unknown. By investigating *gjd4*/Cx46.8, our study provides valuable insights for understanding the functional role of individual Connexins in coordinating critical bioelectric information during development.

## Materials and Methods

### Zebrafish

Fish were maintained by the University of Oregon Zebrafish Facility using standard husbandry techniques (Westerfield, 2000). Animal use protocol AUP-21-42 was approved by the University of Oregon IACUC committee and animal work was overseen by Dr. Kathy Snell.

### scRNAseq

The scRNAseq dataset used in this study originated from the Farnsworth et al., 2020 paper^50^ and was subsequently re-aligned and analyzed in the Lukowicz-Bedford et al., 2022 publication^15^. Larvae samples were pooled, and two replicates were collected at each sampled timepoint (1, 2, and 5 dpf). Standard protocols were employed to dissociate cells from entire larvae^50^. The dissociated cells were then processed using the 10X Chromium platform with 10x v.2 chemistry, aiming to capture approximately 10,000 cells per run. To ensure accurate annotation of *connexin* genes, we examined pooled, full-length RNA-seq and extended the 3’ UTR regions within the gene model (GTF) as necessary^15^. The aligned reads were mapped to the zebrafish genome, GRCz11, using the 10X Cellranger pipeline (version 3.1), utilizing the updated GTF file. For access to the updated GTF file and other related materials, please visit https://www.adammillerlab.com/.

In the analysis, the previously annotated skeletal muscle clusters were first extracted from the large dataset. These clusters were then validated using myogenic markers, including *myod1* and *chrna1*^51–53^. The distinction between slow and fast skeletal muscle clusters was based on the differential expression of canonical markers (Supp. Table 1). Specifically, fast muscle clusters were characterized by elevated expression of *mylpfa*, *myhz2*, while slow muscle clusters were identified by differential expression of *smyhc1*, and *tnnc1b*^51–53^.

### Fluorescent RNA in-situ and immunohistochemistry

To visualize *gjd4* and *gja2* RNA expression, custom RNAscope probes were designed and ordered through ACD (https://acdbio.com/; Supp. Table 2 for details). For fluorescent RNA in situ hybridization, we utilized a modified RNAscope protocol ^86^. Initially, embryos were fixed for 2 hours at room temperature using 4% paraformaldehyde (PFA) and subsequently stored overnight at −20°C in 100% methanol. The fixed tissue was then subjected to protease treatment for 30 minutes, followed by washes with PBS containing 1% Triton X (PBSTx). Next, the tissue was hybridized with the 1× probe overnight at 40°C. Standard RNAscope V2 multiplex reagents and Opal fluorophores were used, with the modification that PBSTx was utilized for all wash steps. The stained tissue was either mounted as whole mounts or cryo-sectioned immediately and mounted with ProLong Gold Antifade (ThermoFisher).

For immunohistochemistry, animals were fixed in 4% paraformaldehyde (PFA) for 2 hours at room temperature, followed by rinsing with PBSTx. The tissue was dehydrated using a methanol series and stored overnight at −20°C in 100% methanol. Subsequently, the tissue was rehydrated through a methanol series and incubated in western block at room temperature for 1 hour. The fish were then exposed to primary antibodies overnight at room temperature (Supp. Table 2 for details). On the following day, the fish were washed with PBSTx at room temperature and incubated with secondary antibodies overnight. After washing, the tissue was subjected to a series of glycerol solutions and then mounted using Prolong Antifade Mount for imaging. For cross sections, tissue was sectioned after staining (either RNA in-situ or immunohistochemistry) on a cryosection, ∼16μm sections.

### Transgenic and mutant line generation

To disrupt *gjd4*, we screened for CRISPR sites using CRISPRScan^87^ and identified sites that would induce mutations at the beginning of the second exon, the primary protein coding region. Single cell embryos were injected with a solution of Cas9 protein and *gjd4*-specific sgRNA (Supp. Table 2 for details)^88^. A subset of the F0 injected embryos were examined for CRISPR-induced genome cutting via PCR. In successfully injected clutches, sibling embryos were raised, and subsequently outcrossed to identify fish with germline transmission of mutations. The *gjd4^b1458^* line was identified as an 8 base pair deletion, resulting in a frame shift in the coding sequence and early stop codon, and was maintained for subsequent analysis (Supp. Table 2 for details).

To tag *gjd4/*Cx46.8, the V5 CRISPR-site was carefully selected based on the predicted Connexin protein structure and using previous approaches that tagged Connexin and did not disrupt localization of function^59,60^. A custom single-stranded oligo was generated with flanking *gjd4*-homology arms, glycine linkers, and the V5-epitope sequence. Single cell embryos were injected with a solution of Cas9 protein, V5-single stranded repair oligo, and *gjd4*-specific sgRNA (Supp. Table 2 for details). A subset of the F0 generation of injected embryos were examined for (1) CRISPR-induced genome cutting via PCR, (2) V5-oligo integration via PCR, and (3) mosaic *gjd4*/Cx46.8-V5 expression in muscle. In successfully injected clutches, sibling embryos were raised, and subsequent crosses were performed to identify germline transmission. The F1 embryos resulting from these crosses were screened both through PCR analysis and immunostaining. Once a founder fish carrying the desired CRISPR-edited *gjd4-v5^b1459^* line was identified, it was outcrossed to ABC-wildtype animals and maintained for analysis (Supp. Table 2 for details).

The *smyhc1:GCaMP3 ^b1464^* line was generated using the plasmid construct described by Jackson et al., 2015^53^. Single-cell embryos were injected with the purified plasmid along with the ISce-1 meganuclease to facilitate integration^53,89^. The F0 generation of injected embryos was raised, and outcrosses were performed to identify fish with germline transmission. Once a founder fish carrying the *smyhc1:GCaMP3* was identified, F1 progeny were screened using immunostaining with specific antibody markers for slow muscle, such as F59 (Fig. S2A). A single *smyhc1:GCaMP3* positive animals was outcrossed to ABC to establish the *smyhc1:GCaMP3 ^b1464^* line. This line was subsequently crossed into the *gjd4^b1458^* mutant background to create the desired experimental line, which was then maintained for further investigations.

### Muscle dye fills and analysis

Glass pipette needles were filled with a 5% solution of sulforhodamine-B^46^ diluted in 0.2M KCl. Fish were anesthetized with MESAB and mounted with 1.2% agar. Single muscle cells were loaded with dye at ∼26hpf and imaged 15 min after each single muscle cell was labeled on an SP8 confocal microscope. To quantify this dataset, an ROI was drawn around the loaded cell, and all surrounding cells in FIJI. The mean fluorescence value was extracted and normalized to the loaded cell. Cells were counted as having dye if they had a value equal to or greater than 5% normalized fluorescence.

### Coiling behavior analysis

To assess the coiling frequency, 22-23hpf embryos were imaged within their chorion, capturing frames at a 30 fps for a duration of 30 seconds on the Ramona Eagle. Regions of interest (ROIs) were manually drawn around the embryos, and the movement was automatically calculated using the Zebrafish STM package in MATLAB^90^. For analyzing the coiling kinematics, embryos were positioned on their back and embedded in 1.2% agar mixed with embryo medium and imaged at 30fp/s for 30 seconds. The angles of tail movements were quantified using FIJI software.

### Calcium imaging and analysis

To examine calcium dynamics, 22-23 hpf *smyhc1:GCaMP3* embryos were mounted on a coverslip in 1.3% agar and embryo medium. Imaging was performed using an SP8 confocal microscope equipped with a 488 nm laser. Z-stacks with a thickness of 7 micrometers were acquired at a frame size of 256×256 and an imaging speed of 700Hz, resulting in a total stack time of approximately 2.13 seconds per stack for a total of 330seconds. Subsequently, maximum intensity projections of 3 micrometer sections were generated. To correct for any motion artifacts during the imaging process, the moco package in FIJI^91^ was utilized, employing the first frame of each stack for reference. In FIJI, regions of interest (ROIs) were carefully hand-drawn around individual cells to ensure accurate capture of the cell’s fluorescence signal. Mean fluorescence values from the ROIs were extracted, and the relative change in fluorescence (Δf/F) was calculated by taking the average fluorescent value from the 5-10th percentile of raw fluorescence values across the entire recording period.

Correlation of cellular activity throughout the recording period was calculated using a Pearson correlation - where r was computed for every pair of cells over recording time with 95% confidence intervals and plotted in a heatmap matrix. Cross-correlation was calculated by using the ‘Xcorr’ function in MATLAB. In brief, f/F traces for recording sessions were imported, and Xcorr was run between all cells in the same animal. To quantify the average and maximum Δf/F for each animal, the Δf/F values across cells in a recording period were used. The average Δf/F was determined by computing the mean of all Δf/F values for the cells in an animal, while the maximum Δf/F was determined by identifying the highest Δf/F value across the cells in an animal. The firing rate in Hz was computed by calculating the standard deviation of the 1st to 5th percentile of each cell’s Δf/F values from the entire recording period. This standard deviation was multiplied by 15 to establish a cell-specific threshold, which was used as input for the ‘FindPeaks’ function in MATLAB. This process was conducted for each cell during the first minute of recording. The Hz firing rate for each cell was then averaged across the cells within an animal.

### Gap junction blockers

To block gap junction channels, we used three different compounds previously shown to interfere with channel transmission including heptanol, octanol and quinine^46,62,72,73^. We exposed embryonic fish (22-24hpf) to differing concentrations of this compounds in embryo medium (EM) and screened for a halt of coiling behavior. We found that 100um of heptanol, 200um of octanol, and 0.25M of quinine were sufficient to stop coiling. We found that coiling stopped with ∼10 min of application to 100um of heptanol and 200um octanol, and within ∼25 min of application of quinine. Once fish were transferred to fresh EM, coiling returned within ∼10min for heptanol and octanol, and ∼25 min for quinine. For calcium imaging, fish were de-chorionated and allowed to soak in the GJ blocker solutions for times specified above before imaging. The success of pharmacological manipulations was assessed by screening the fish for no movement. For myosin imaging, fish were exposed to the corresponding drugs, except for quinine, at 16hpf and were euthanized and fixed at 24hpf. Fish exposed to the quinine solution at 16hpf did not survive to 24hpf.

### Bungarotoxin injections

A solution of α-bungarotoxin (α-btx) at a concentration of 1mg/mL was injected into the yolk of approximately 16-17 hours post-fertilization (hpf) zebrafish embryos at the pre-coiling stage^75^. Subsequently, the injected animals were raised under regular conditions until they reached 23-24 hpf for analyses. The success of α-btx injections was assessed by screening the fish for no movement. For comparison, sham injections were also performed using a similar procedure and conditions. However, instead of α-btx, the sham injections involved the use of ddH20, which served as the diluent for the α-btx solution.

### Myosin structure analysis

To detect slow muscle myosin, we used the F59 antibody, which specifically recognizes myosin heavy chains, following standard immunohistochemistry protocol^47^. The fish samples were imaged using an SP8 confocal microscope. To quantify the tortuosity of the myosin fibers, a straight line was drawn across the length of the somite in FIJI. Subsequently, another line was drawn through the center of the myosin fiber. The tortuosity was determined by calculating the ratio of the somite length over the true length of the myosin fiber^78^. For each cell type within an animal, five tortuosity measurements were taken across each cell type, and the reported values represent the average of these measurements. For sarcomere length measurements, a straight line was drawn across the length of the sarcomere using FIJI. We recorded ten sarcomere measurements per cell type within each animal, and report values that represent the average of these measurements.

### Muscle cell fate specification and somite structure analysis

To quantify muscle cell types we used antibody staining of the engrailed and prox1 transcription factors (see Supp. Table 2 for antibody details). Engrailed alone marks the medial fast fibers (MFF), prox1 alone marks the superficial slow fibers (SSF) and co-expression of prox1 and engrailed marks the muscle pioneers (MP)^77^. To calculate the number of cells per somite, cell nuclei were counted in 3 somites over the yolk extension (somites ∼12 through 15) and counts/somite were averaged for each animal.

To examine the structure of the somite and other features of the muscle, we used the anti-laminin, anti-dystrophin, and anti-myosin heavy chain (MF20) antibodies (see Supp. Table 2 for antibody details)^46^. Approximately 3 somites over the posterior of the yolk extension (somites ∼15-18) were imaged.

## Acknowledgments

We would like to thank the entire Miller lab for their comments, suggestions, and guidance throughout experiments and the drafting of this manuscript. We thank Anisha Adke, who contributed to the generation and maintenance of the *gjd4^b1458^* line and Dr. Sarah Stednitz for initial characterization of the behavioral phenotype of these animals. We thank Dr. Yosuke Ono who provided us with the *smyhc1:GCaMP3* plasmid, Dr. Amy Robbins who provided us with the *ptch2:kaede* line, and Ellie Melancon and Jeremy Wagner for many antibodies. We thank Dr. PK Loi and the UO histology team for sectioning tissue, and the entire UO Fish facility for the exceptional care of our animals. We thank Dr. Matt Smear for his advice with calcium data analysis.

## Author Contributions

RML-B performed experiments and analysis, except for the dye fills that were performed by JSE. RML-B and ACM wrote the manuscript, with editing from JSE. ACM acquired funding.

## Supplementary Materials

**Figure S1:**
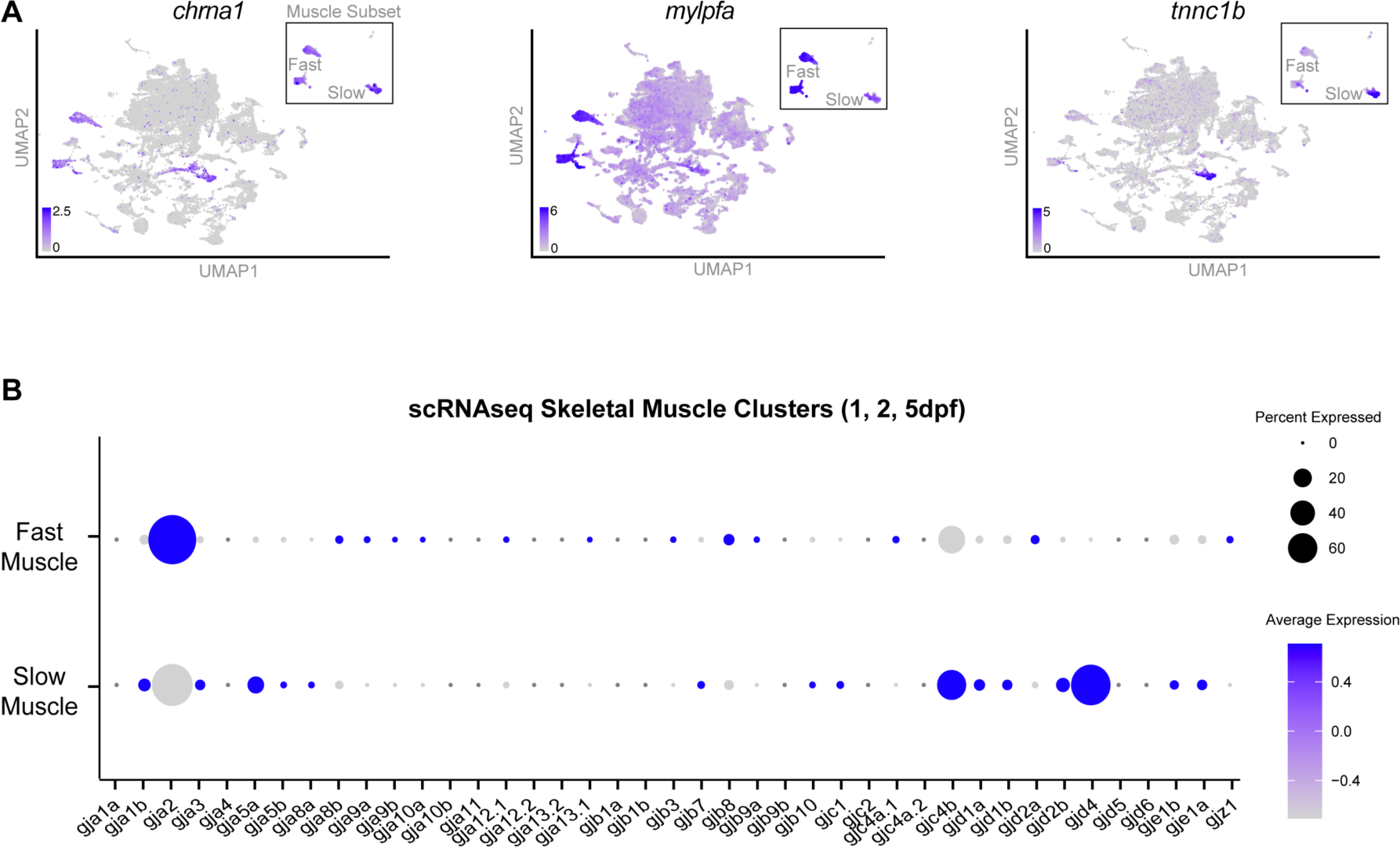
*connexin* expression in the skeletal muscle. **A)** Expression of skeletal muscle markers *chrna1* (all muscle), *mylpfa* (enriched in fast subtypes), and *tnnc1b* (slow subtypes), are shown for all cells of the scRNAseq atlas dataset. Expression within only the putative skeletal muscle clusters is shown as subsets. **B)** Expression of the entire zebrafish *connexin* gene family across the putative fast or slow skeletal muscle clusters. Connexin genes are arranged alphabetically, and expression is denoted by the size of the circle and intensity of the color.

**Figure S2:**
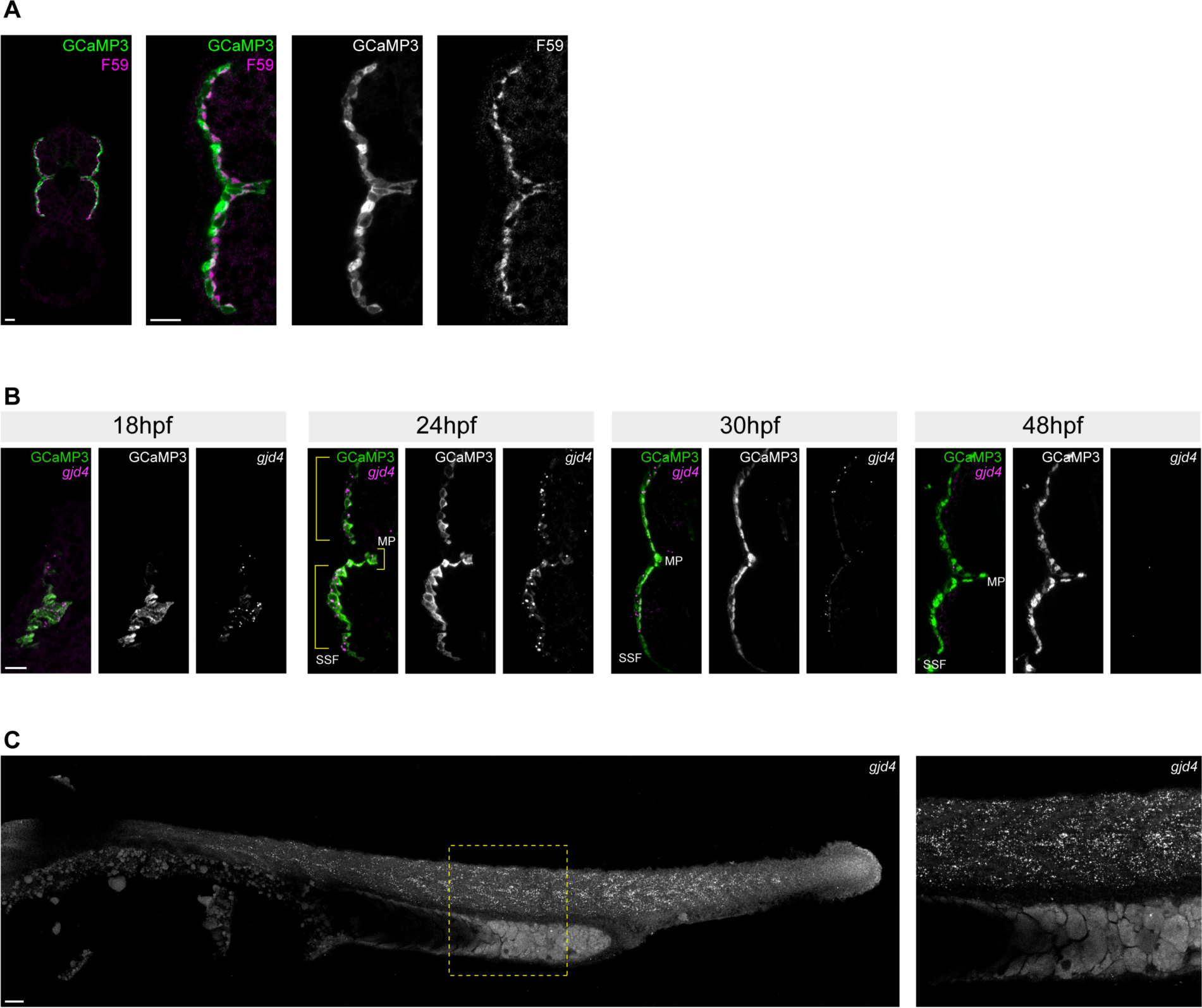
Validation of *smyhc1:GCaMP3^b1464^* and *gjd4* expression. **A)** Cross-section of *smyhc1:GCaMP3^b1464^* line (green) with the slow muscle marker F59 (magenta) at 24hpf. Scale bar = 10μm. **B)** Fluorescent RNA in situ hybridization of *gjd4* at specified times (hours post fertilization (hpf)). Cross sections through the mid-trunk of embryos, where slow muscle cells are marked by the *smych1:GCaMP^b1464^* line, shown in green, and *gjd4* transcripts are shown in magenta. Yellow brackets highlight SSF cells and MP cells. Scale bar = 10μm. **C)** Whole mount RNA in-situ against *gjd4* in a 24hpf embryo. Yellow box indicates a zoom shown in right. Scale bar = 25μm.

**Figure S3:**
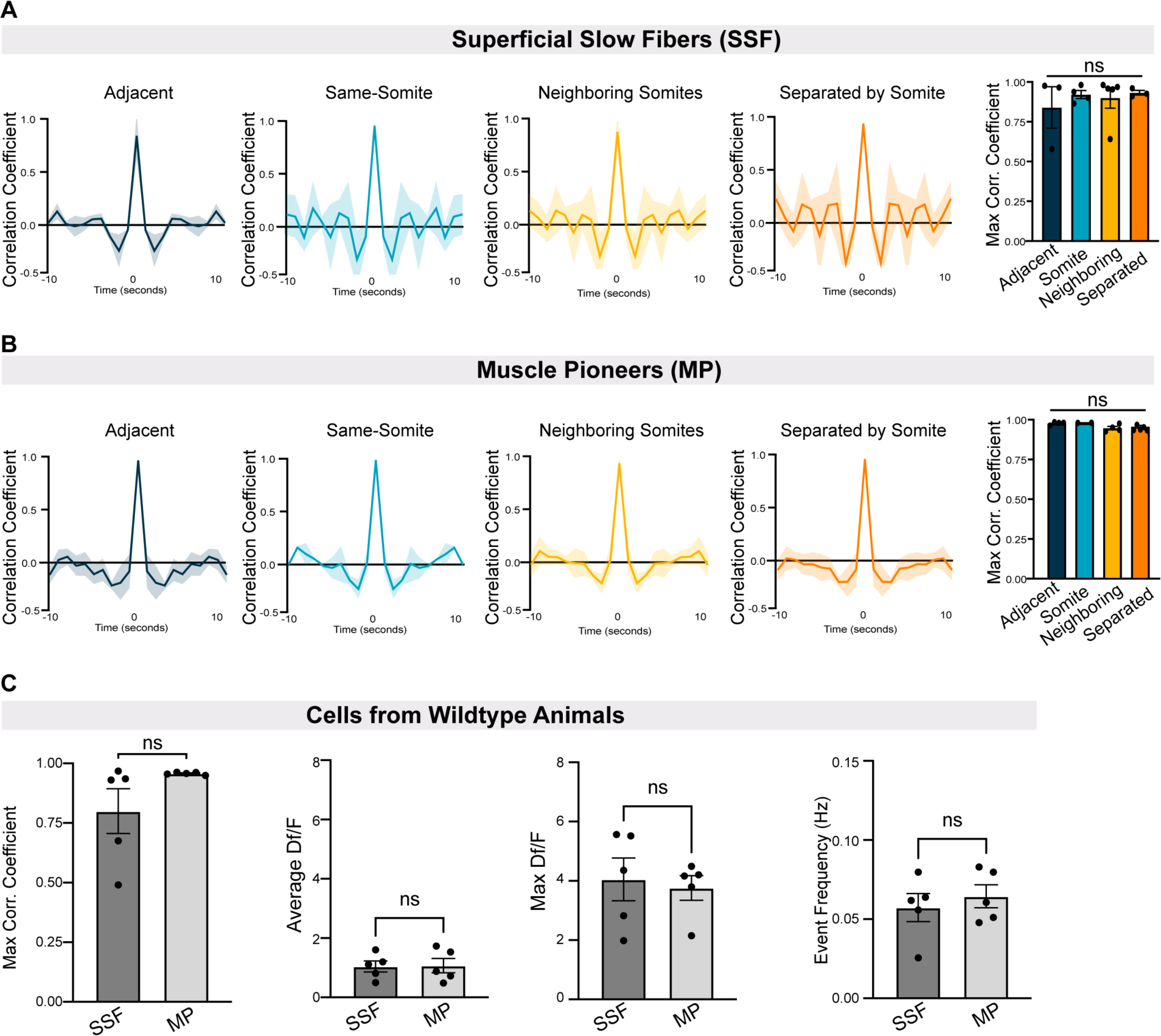
Dynamics of calcium activity in the SSF and MP cells. **A-B)** Cross correlation analysis relative to cellular location. **A)** Cross-correlation analysis of SSF cells in wildtype showing mean (dark line) and standard deviation (shading) of adjacent cells (dark blue), cells from the same somite (light blue), cells from neighboring somite (yellow), and cells separated by one somite (orange). Max correlation coefficient for each classification shown at right, Brown-Forsythe ANOVA test with multiple comparisons, adjacent vs. somite p-value = 0.9753, adjacent vs. neighbor p-value = 0.9967, adjacent vs. separate p-value = 0.9565, somite vs. neighbor p-value = 0.997, somite vs. separated p-value = 0.9983, neighbor vs. somite p-value = 0.9946. **B)** Cross-correlation analysis of MP cells in wildtype showing mean (dark line) and standard deviation (shading) of adjacent cells (dark blue), cells from the same somite (light blue), cells from neighboring somite (yellow) and cells separated by one somite (orange). Max correlation coefficient for each classification shown at right, Brown-Forsythe ANOVA test with multiple comparisons, adjacent vs. somite p-value = 0.9673, adjacent vs. neighbor p-value = 0.2500, adjacent vs. separate p-value = 0.2277, somite vs. neighbor p-value = 0.1939, somite vs. separated p-value = 0.1145, neighbor vs. somite p-value = 0.9467. **C)** Comparison of GCaMP fluorescence activity between WT SSF and MP cells. N = 5 per cell type. Unpaired t-test with Welch’s correction for each, max correlation coefficient p-value = 0.1668, average Δf/F p-value = 0.9334, max Δf/F p-value = 0.7400, and event frequency p-value = 0.5535.

**Figure S4:**
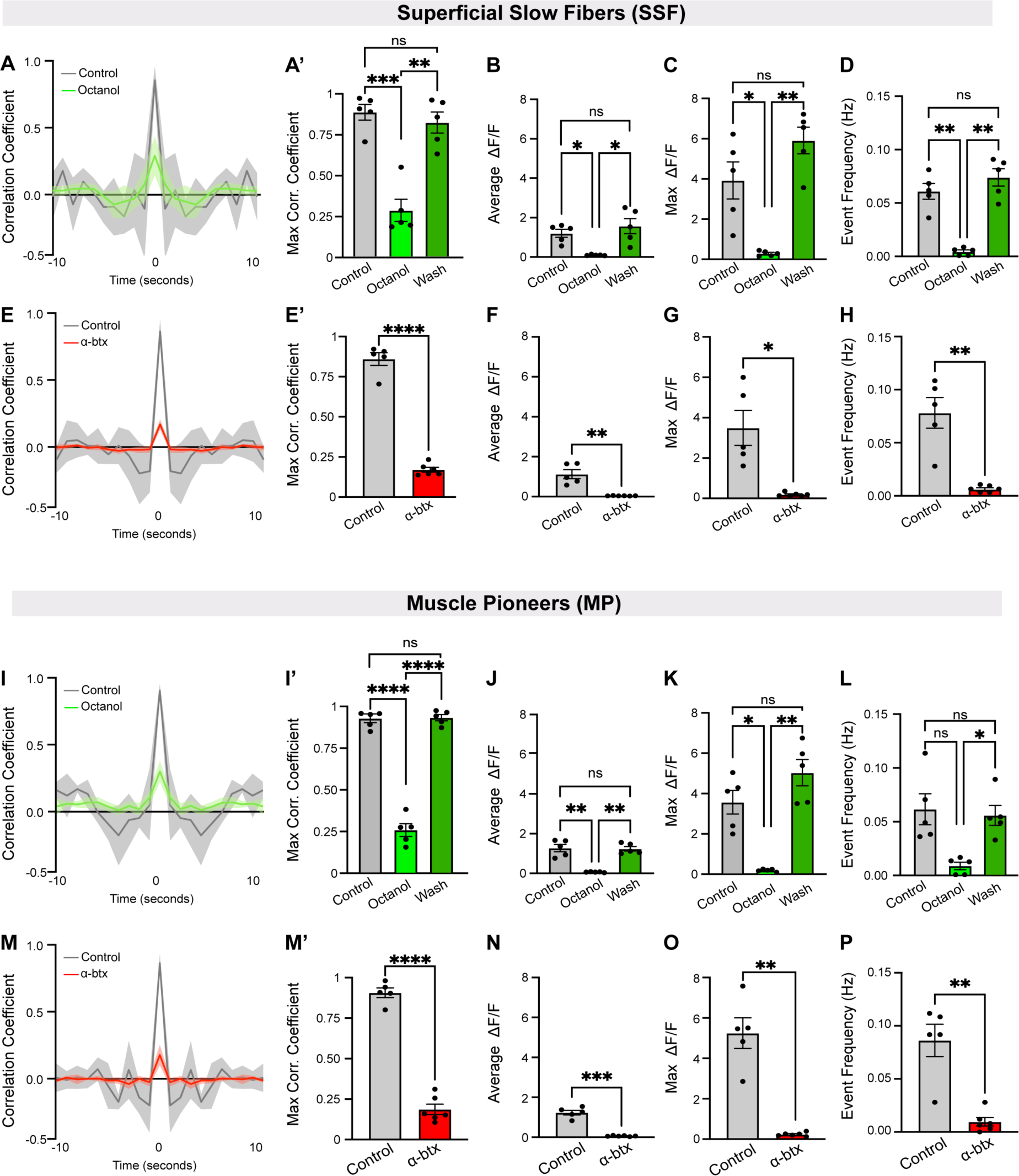
Spontaneous calcium activity in slow muscle cells with octanol or alpha-bungarotoxin treatment. **A-H)** Analysis of calcium dynamics in SSF cells. **A)** Cross-correlation analysis of SSF cells showing mean (dark line) and standard deviation (shading) of control (grey) and octanol treated (green) embryos. N = 5 animals per condition. Octanol-wash cross-correlation traces were omitted for graphical clarity. **A’)** Max correlation coefficient for each animal, Brown-Forsythe ANOVA test with multiple comparisons, control vs. control vs. octanol p-value = 0.0005, control vs. wash p-value = 0.8151, octanol vs. wash p-value = 0.0013. **B)** Average GCaMP fluorescence across the recording session for each animal. N = 5 per condition, Brown-Forsythe ANOVA with multiple comparisons, control vs. octanol p-value = 0.0143, control vs. wash p-value = 0.7844, and octanol vs. wash p-value = 0.0463. **C)** Max GCaMP fluorescence across the recording session for each animal. N = 5 per condition, Brown-Forsythe ANOVA test with multiple comparisons, control vs. octanol p-value = 0.0430, control vs. wash p-value = 0.3018, and octanol vs. wash p-value = 0.0028. **D)** Average peak frequency across the recording session for each animal. N = 5 animals per condition, Brown-Forsythe ANOVA test with multiple comparisons, control vs. octanol p-value = 0.0044, control vs. wash p-value = 0.5791 and octanol vs. wash p-value = 0.0028. **E)** Cross-correlation analysis of SSF cells showing mean (dark line) and standard deviation (shading) of control (grey) and α-btx injected (red) embryos. N = 5 control, 6 α-btx injected. **E’)** Max correlation coefficient for each animal, unpaired t-test with Welch’s correction, p-value = <0.0001. **F)** Average GCaMP fluorescence across the recording session for each animal. N = 5 control, 6 α-btx injected, unpaired t-test with Welch’s correction, p-value = 0.0086. **G)** Max GCaMP fluorescence across the recording session for each animal. N = 5 control, 6 α-btx injected, unpaired t-test with Welch’s correction, p-value = 0.0189. **H)** Average peak frequency across the recording session for each animal. N = 5 control, 6 α-btx injected, unpaired t-test with Welch’s correction, p-value = 0.0072. I-L) Analysis of calcium dynamics in MP cells. **I)** Cross-correlation analysis of MP cells showing mean (dark line) and standard deviation (shading) of control (grey) and octanol treated (green) embryos. N = 5 animals per condition. Octanol-wash cross-correlation traces were omitted for graphical clarity. **I’)** Max correlation coefficient for each animal, Brown-Forsythe ANOVA test with multiple comparisons, control vs. octanol p-value = <0.0001, control vs. wash p-value = 0.9992, octanol vs. wash p-value = <0.0001. **J)** Average GCaMP fluorescence across the recording session for each animal. N = 5 per condition, Brown-Forsythe ANOVA test with multiple comparisons, control vs. octanol p-value = 0.0070, control vs. wash p-value = 0.9982, and octanol vs. wash p-value = 0.0014. **K)** Max GCaMP fluorescence across the recording session for each animal. N = 5 per condition, Brown-Forsythe ANOVA test with multiple comparisons, control vs. octanol p-value = 0.0115, control vs. wash p-value = 0.3215, and octanol vs. wash p-value = 0.0045. **L)** Average peak frequency across the recording session for each animal. N = 5 per condition, Brown-Forsythe ANOVA test with multiple comparisons, control vs. octanol p-value = 0.0584, control vs. wash p-value = 0.9806 and octanol vs. wash p-value = 0.0134. M-P) Analysis of calcium dynamics in α-btx MP cells. **M)** Cross-correlation analysis of MP cells showing mean (dark line) and standard deviation (shading) of control (grey) and α-btx injected (red) embryos. N = 5 control, 6 α-btx injected. **M’)** Max correlation coefficient for each animal, unpaired t-test with Welch’s correction, p-value = <0.0001. **N)** Average GCaMP fluorescence across the recording session for each animal. N = 5 control, 6 α-btx injected, unpaired t-test with Welch’s correction, p-value = 0.0006. **O)** Max GCaMP fluorescence across the recording session for each animal. N = 5 control, 6 α-btx injected, unpaired t-test with Welch’s correction, p-value = 0.0027. **P)** Average peak frequency across the recording session for each animal. N = 5 control, 6 α-btx injected, unpaired t-test with Welch’s correction, p-value = 0.0058.

**Figure S5:**
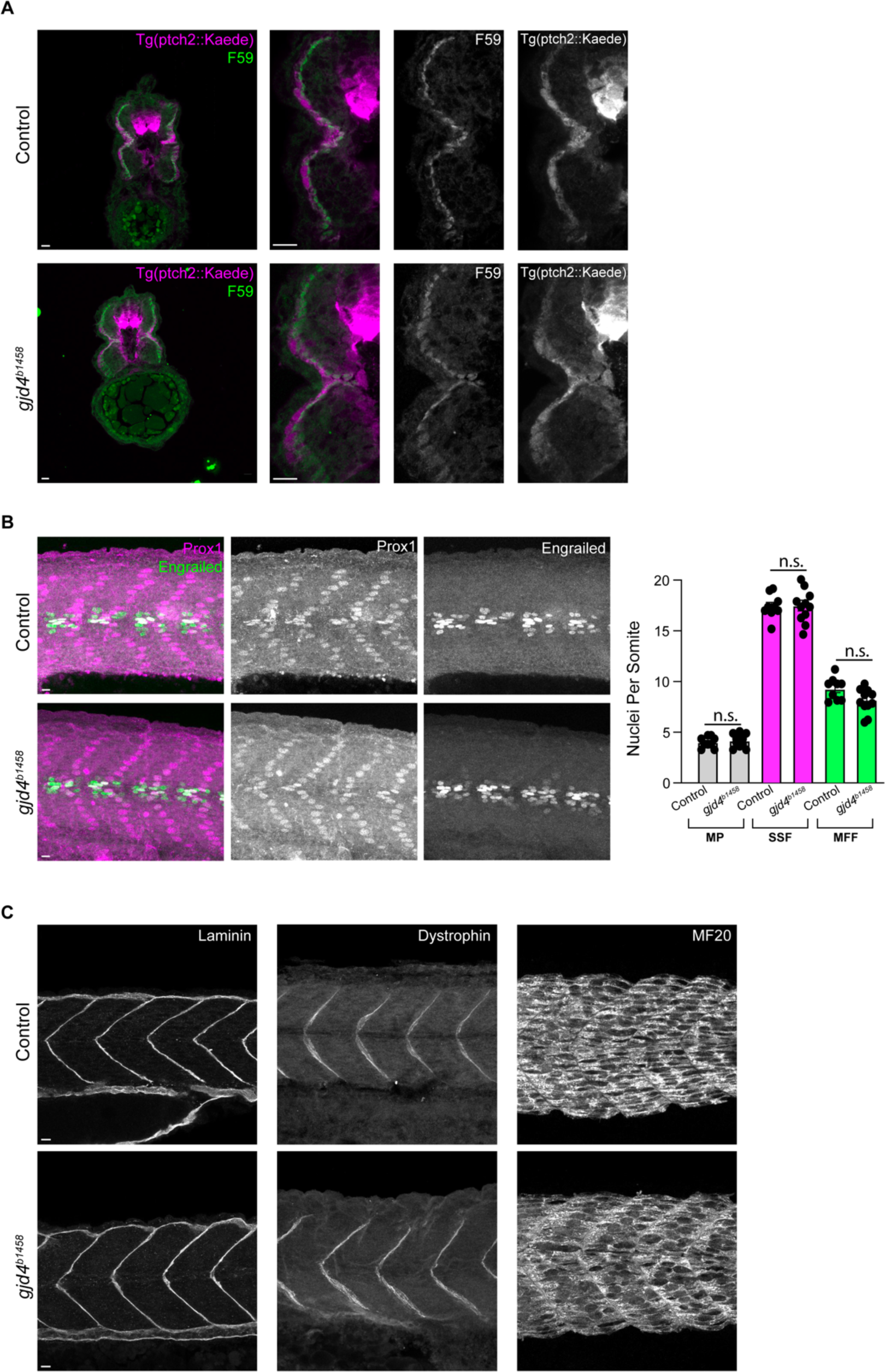
Loss of *gjd4*/Cx46.8 does not result in overall patterning defects. **A)** Representative images from cross-section of control *ptch2:kaede ^a4596^* and *ptch2:kaede ^a4596^*; *gjd4^b1458^* mutants, stained with the slow muscle marker F59 (green). Scale bar = 10μm. **B)** Counts of skeletal muscle cell types using prox1a, which alone marks SSF cells, engrailed, which alone marks the medial fast fiber (MFF) cells, and co-expression of prox1a and engrailed which marks the MP cells. Cell nuclei were counted in 3 somites per animal and averaged for each animal. N = 9 control, and 11 *gjd4^b1458^* mutants, unpaired t-test with Welch’s correction between genotypes for each cell type, MP p-value = 0.5536, SSF p-value = 0.8971 and MFF p-value = 0.0512. **C)** Representative images for different muscle structures, including laminin, dystrophin, and fast muscle (MF20), at 24hpf in control and *gjd4^b1458^* mutant. Scale bar = 10μm.

**Figure S6:**
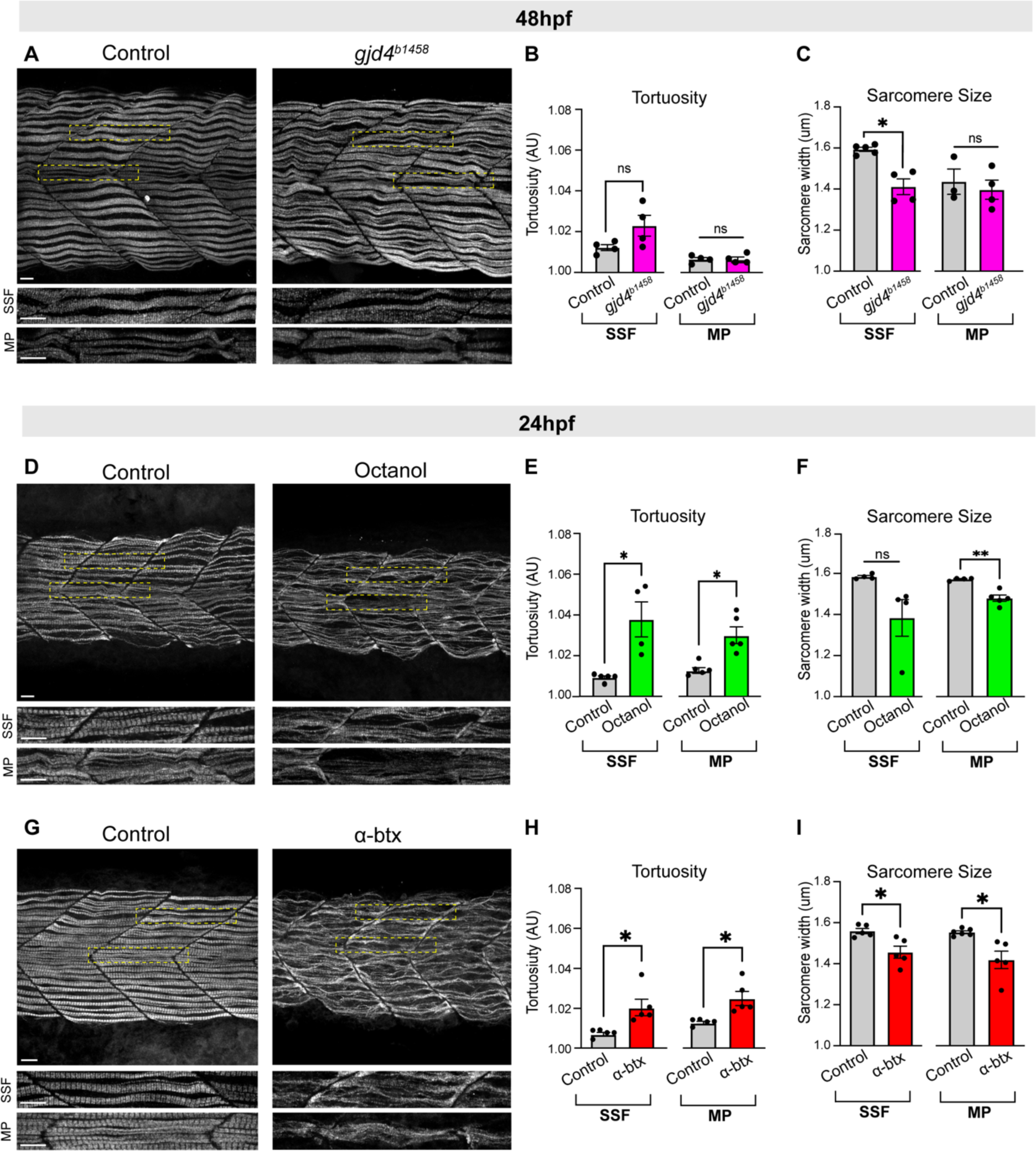
Disruption to GJ communication results in disrupted slow myosin fibers. **A)** Representative image of slow muscle myosin fibers at 48 hpf in control and *gjd4^b1458^* mutant animals. Yellow-dashed boxes indicate zoomed regions for the SSF or MP, shown in neighboring panels below. Scale bar = 10μm. **B)** Quantification of tortuosity. N = 4 animals per genotype, unpaired t test with Welch’s correction between each genotype, for each cell type, SSF p-value = 0. 1248, and MP p-value = 0. 8929. **C)** Quantification of sarcomere size. N = 4 animals per genotype, unpaired t test with Welch’s correction between each genotype, for each cell type, SSF p-value = 0.0142, and MP p-value = 0.6020. **D)** Representative image of slow muscle myosin fibers at 24 hpf in control or animals exposed to octanol. Yellow-dashed boxes indicate zoomed regions for the SSF or MP, shown in neighboring panels below. Scale bar = 10μm. **E)** Quantification of tortuosity. N = 5 SSF control, 4 SSF octanol, and 4 animals per condition for MP. Unpaired t test with Welch’s correction between each condition, for each cell type, SSF p-value = 0. 1248, and MP p-value = 0. 8929. **F)** Quantification of sarcomere size. N= 4 control SSF and 5 octanol SSF, N = 4 animals per condition for MP. Unpaired t test with Welch’s correction between each condition, for each cell type, SSF p-value = 0.113, and MP p-value = 0.0032. **G)** Representative image of slow muscle myosin fibers at 24 hpf in control or α-btx injected animals. Yellow-dashed boxes indicate zoomed regions for the SSF or MP, shown in neighboring panels below. Scale bar = 10μm. **H)** Quantification of tortuosity. N = 5 animals per condition. Unpaired t-test with Welch’s correction between each genotype, for each cell type, SSF p-value = 0.0294, and MP p-value = 0.0247. **I)** Quantification of sarcomere size. N = 5 animals per condition. Unpaired t test with Welch’s correction between each genotype, for each cell type, SSF p-value = 0.0207, and MP p-value = 0.0329.

